# Mitochondrial Fatty Acid Oxidation is Stimulated by Red Light Irradiation

**DOI:** 10.1101/2024.09.12.612633

**Authors:** Manuel Alejandro Herrera, Camille C. Caldeira da Silva, Mauricio S. Baptista, Alicia J. Kowaltowski

## Abstract

Keratinocytes are the primary constituents of sunlight-exposed epidermis. In these cells, UVA completely inhibited oxidative phosphorylation, while equivalent doses of blue and green light preserved metabolic fluxes, but reduced viability. In contrast, red light enhanced proliferation and elevated basal and maximal oxygen consumption rates for 48 h, without altering protein levels of the electron transport chain. Targeted flux analysis revealed that red light specifically activates AMPK-dependent mitochondrial fatty acid oxidation. This was accompanied by reduced the levels of free fatty acids and increased acetyl CoA carboxylase phosphorylation. Together, our results characterize wavelength-selective regulation of keratinocyte metabolism: UV/visible wavelengths induce damage, while red light triggers AMPK-dependent fatty acid oxidation, providing a mechanistic explanation for photobiomodulation in epidermal cells.

**Highlights:** UVA (365 nm) abolishes oxidative phosphorylation, while blue/green light (450/517 nm) induce photosensitized toxicity, without metabolic disruption.

Red light (660 nm) boosts keratinocyte proliferation and sustains elevated mitochondrial respiration for 48 h.

Permeabilized cell electron transport remains unchanged, implicating cytosolic (not mitochondrial) regulation of fuel utilization.

Red light activates AMPK and phosphorylates/inhibits acetyl CoA carboxylase (ACC).

Photobiomodulation with red light occurs through AMPK/ACC-mediated lipid oxidation, not direct respiratory chain modulation.

## 1. Introduction

Living organisms evolved mechanisms to sense and respond to environmental stimuli [1], including light [2, 3], and specifically wavelengths emitted from the sun in the ultraviolet (UV, 320 to 400 nm) and visible light (400 to 700 nm) ranges [4]. Energy from the photons in both the UV and visible ranges can interact with different biomolecules, scattering, transmitting, reflecting, or absorbing light according to the optical properties of the components of that specific organism. The most recognized consequence of photon absorption is the generation of excited states, which are much more reactive than their ground states [4] and can interact with surrounding molecules, exciting other molecules and promoting chain reactions that may oxidize others (photosensitized oxidation) [5], a common undesirable effect of light.

The tissue most exposed to solar radiation in the human body is the skin, which is also our largest organ, vital in the maintenance of our body, offering an effective barrier against external agents, among many other functions. The skin involves a large number of cell types with different functions and relative abundance depending on the skin layer; in the epidermis, the most exposed layer to radiation, keratinocytes are the most common [6]. Indeed, endogenous photosensitizing molecules in keratinocytes such as flavins, nicotinamides, and cytochromes, located mostly in mitochondria, can produce excited and reactive species, affecting keratinocyte and skin function in response to light [7].

Interestingly, light exposure also has benefits to humans. Several light-based technologies are used to treat a variety of diseases, including photobiomodulation, in which light at different wavelengths is used to irradiate human skin, causing several beneficial effects, including pain control and wound healing, that can be useful in the treatment of inflammatory, neurological, and musculoskeletal disorders [8]. Although the molecular mechanisms responsible for photobiomodulation are not completely understood, mitochondrial functions such as oxidative phosphorylation are often found to play an important role [9].

Indeed, mitochondria are pivotal for cell homeostasis as the center for energy metabolism, as well as having roles in cell growth, adaptation, death, ion homeostasis, oxidant production, and thermogenesis, among others [10]. Many of these functions, and especially the canonic function of oxidative phosphorylation, require endophotosensitizers such as NADH, FADH_2_, and metal-containing proteins and cytochromes in respiratory complexes I, III, and IV [11]. As a result, mitochondria and oxidative phosphorylation are an expected target of cellular functional consequences of light exposure.

While the role of keratinocytes in light-induced melanogenesis via non-neuronal pathways is well-documented [12], light-dependent effects on keratinocyte bioenergetics remain unexplored. Using metabolic flux analysis, we demonstrate here that specific light wavelengths differentially modulate oxidative phosphorylation and cellular integrity, with UV/visible spectra inducing photodamage, while red light (660 nm) enhances mitochondrial respiration. Remarkably, we identify a novel mechanism underlying red light effects: selective activation of fatty acid oxidation via AMPK-mediated metabolic reprogramming, independent of direct electron transport chain modulation. These findings establish keratinocytes as wavelength-sensitive metabolic sensors and redefine photobiomodulation paradigms in the epidermis.

## 2. Material and Methods

### 2.1. Cell Cultures

A human immortalized keratinocyte cell line (HaCaT, [13]), an immortalized skin fibroblast cell line (Hs68), and a melanocyte cell line (B16F10) were cultured in high glucose Dulbecco modified eagle medium (DMEM) with phenol red (Gibco, Life Technologies, USA), supplemented with 10% v/v fetal bovine serum (FBS; Sigma), 110 mg/mL sodium pyruvate, 4 mM L-glutamine, 100 U/mL of penicillin, and 100 pg/mL streptomycin (Gibco, Life Technologies, USA) at pH 7.4, 37°C in a humidified atmosphere of 5% CO_2_. Passages were kept between 20 and 30 for all cell lines, using 0.25% trypsin-EDTA for enzymatic dissociation. Cells were counted manually in a hemocytometer in the presence of trypan blue. Notably, primary cells undergo major metabolic alterations during and after isolation [14], justifying the use of cell lines.

### 2.2. Irradiation Sources and Conditions

Irradiation conditions for different wavelengths were adjusted by dose to a final 36 J/cm^2^. For UVA, this was achieved at 365 nm λmax (light source from Biolambda, Brazil), for blue light at 450 nm λmax (light source from Biolambda, Brazil), for green light with at 517 nm λmax (light source from Biolambda, Brazil), and for red light with at 660 nm λmax (light source from Ethik Technology, Brazil). Characterization of the emission sources was performed by acquiring emission spectra using a USB2000 + UV-VIS-ES spectrometer and a QP50–2-UV-VIS (Ocean Optics, USA) optical fiber. Irradiance was measured using a Laser Power Meter (Coherent Inc., Santa Clara, CA, USA) containing a silicon power sensor for the visible range (LM-2 Vis, 1,061,323); LED spectrum emissions are presented in Supplementary Figure S1, and energy specifications and number of photos are presented in Table 1 and Table 2 of the supplementary material. All cells were plated 24 hours before the experiment. Prior to irradiation treatment, they were washed and incubated in phosphate buffer saline (PBS), in an irradiation chamber without CO_2_, and with controlled temperatures of 37°C. All treatments were done in PBS and in the chamber for the same time of incubation, including dark controls. Changes in local temperature in wells were determined using a thermal handheld InfiRay P series camera (Hefei, China).

### 2.3. Cell Survival Assays

#### 2.3.1. MTT assay

Cell survival was estimated based on the reduction of 3-(4,5-dimethylthiazol-2-yl)- 2,5-diphenyltetrazolium bromide (MTT). Exponentially growing HaCaT cells were plated (5 × 10^4^ cells/well) in 48 well cell culture plates (Corning Costar) for 24 h. After 2, 48, and 72 hours of the treatment, we added 0.5 mL of medium containing MTT (Sigma, 50 μg/mL) to each well and incubated at 37 °C for 3 h. The medium with MTT was then removed and 0.25 mL dimethyl sulfoxide (DMSO; Sigma) was added. The plate was shaken and absorbance values were read at 550 nm using a microplate reader (Expectramax i3, Molecular Devices, USA) with a wavelength correction set at 550 nm. Backgrounds in wells without cells were determined and subtracted.

#### 2.3.2. Cristal Violet staining

Cell survival was also estimated by the accumulation of crystal violet (CV) in fixed cells. Exponentially-growing HaCaT cells were plated (5×10^4^ cells/well) in 48 well cell culture plates (Corning Costar) for 24 h. After red light treatments (2, 48, and 72 hours), cells were washed twice with PBS, fixed in alcoholic-based 1.0% v/v acetic acid for 10 min, washed twice with PBS and stained with CV (Sigma) at 0.02% w/v for 5 min at room temperature. After staining, cells were washed again with PBS to eliminate the non-ligated CV. CV was then eluted in 0.1 M sodium citrate - 50% v/v ethanol. Absorbance was read using the Espectramax i3 at 585 nm with a wavelength correction set at 585 nm for the subtraction of backgrounds in wells without cells.

#### 2.3.3. Neutral Red incorporation assay

Cell survival was further estimated by accumulation of Neutral Red (NR), which dyes lysosomes in intact viable cells. Exponentially growing HaCaT cells were plated (5×10^4^ cells/well) in 48 well cell culture plates (Corning Costar, cod. 3548) for 24 h. After red light treatment (2, 48, and 72 hours), cells were washed twice with PBS and stained with 30 μg/mL NR (Sigma) at 37°C for 2 h and subsequently washed twice with PBS. NR was eluted with an alcoholic-based 1.0% v/v acetic acid fixing solution for 10 min at room temperature and measured at 540 nm using an Expectramax microplate reader with wavelength correction set at 540 nm for subtraction of backgrounds in wells without cells.

### 2.4. Extracellular Metabolic Flux Analysis

#### 2.4.1. Cell plating and irradiation

HaCaT were seeded at 1.5 x10^5^ cells per well, melanocytes at 1 x10^5^ and fibroblasts were seeded at 1 x10^5^ on Agilent Seahorse XF24 cell culture microplates in 500 μL high glucose DMEM with 10% FBS, 1% Penicillin/Streptomycin, without phenol red, and allowed to adhere overnight. Plates were treated with light in the absence of CO2, after which the cells were returned to the CO2 incubator for 2 hours of recovery. Post-recovery time, cells were washed twice with 500 μL high glucose DMEM containing 1% penicillin/streptomycin. Media did not contain bicarbonate or FBS. Cells were kept for 1 h in a humidified incubator at 37°C without CO2, after which Oxygen Consumption Rates (OCR) were measured using a Seahorse Extracellular Flux Analyzer (Agilent Technologies).

#### 2.4.2. Mito-Stress test

After the preparation phase described above and incubation in bicarbonate-free media, plates were placed in the Seahorse Extracellular Flux Analyzer and OCR were measured under basal conditions, followed by injections of: a) oligomycin (final concentration 1 μM); b) CCCP (final concentration 1 μM); c) rotenone and antimycin (R/AA, 1 μM each final concentration). Unless otherwise specified, measurements were conducted with 3 min mixing + 1 min waiting + 2 min of measurements. Cell-free wells were incubated with the same medium for background correction, calculated by subtracting changes observed from the experiments in the presence of cells. OCR values obtained were normalized per amount of total protein in each well. For that, at the end of each experiment, the medium was removed, cells were PBS-washed, lysed with RIPA buffer, and total protein was quantified using the Pierce BCA Protein Assay kit.

#### 2.4.3. Fatty acid and glutamine-supported OCRs

After the preparation phase described in 2.4.1 and incubation in bicarbonate-free media, Seahorse XF24 plates were placed in the equipment and OCRs were measured under basal conditions, followed injections of: a) etomoxir (final concentration 40 μM) or BPTES (final concentration 7 μM) or the same volume of DMEM in control wells; b) oligomycin (final concentration 1 μM); c) Carbonyl cyanide m-chlorophenyl hydrazone (CCCP, final concentration 1 μM); d) antimycin and rotenone (R/AA, 1 μM final concentration each). Measurements were conducted and normalized as described in 2.4.2.

#### 2.4.4. Glycolysis stress test

Post-recovery from irradiation, cells were washed twice with 500 μL DMEM with no phenol red containing 1% penicillin/streptomycin. Media did not contain bicarbonate nor FBS. Cells were kept for 1 h in a humidified incubator at 37°C without CO_2_. After incubation, Extra-Cellular Acidification Rates (ECARs) were measured using a Seahorse Extracellular Flux Analyzer (Agilent Technologies) under basal conditions, followed injections of: a) glucose (final concentration 25 mM); b) oligomycin (final concentration 1 μM); c) 2-deoxy-glucose (2DG, final concentration 50 mM). Measurements were conducted and normalized as described in 2.4.2.

### 2.5. High Resolution Respirometry

Cells at 80% confluence were treated with red light at 0 J/cm^2^ and 36 J/cm^2^ and submitted to a recovery period as described in 2.4.1. Cells were then detached from the plate using 0.25% trypsin-EDTA, counted in a Countess 3 Cell Counter (Invitrogen, USA) in the presence of trypan blue. Oxygen consumption was measured using a high-resolution Oxygraph-2k (O2k) respirometer (Oroboros Instruments, Austria). One million cells were incubated in 1 mL of experimental buffer (250 mM sucrose, 100 μM EGTA, and 1 mg/mL bovine serum albumin) and different substrates described in the legend at 37°C with continuous stirring (400 rpm) with the light off in both chambers. After initial respiration was measured (state 2), state 3 respiration was measured by adding 1 mM ADP, followed by addition of 1 μM oligomycin (state 4), and 1 μM CCCP (state 3u). Respiratory control ratios (RCRs) were calculated as state 3/state 4.

### 2.6. Long-Term OCR Measurements

To measure metabolic effects of red light long-term, we followed OCRs in plated cells in CO_2_ incubators under normal culture conditions using a Resipher Real-time Cell Analyzer (Lucid Scientific, Atlanta, GA, USA) for 3 days. 1 x 10^5^ cells per well were cultured in a 96 Well black/clear Bottom (Corning Costar) plate using high DMEM without phenol (25 mM glucose, 10% FBS) and allowed to adhere overnight. The cell plates were treated with different doses of red light (0, 6, 12, 36 and 150 J/cm^2^) and the Resipher sensor system was immediately placed above the plate. Measurements of basal OCRs were followed over three consecutive days in culture, in five technical replicates. Cell-free wells were included with the same media for background correction. Media was changed daily.

### 2.7. Western Blots

In order to quantify proteins related to the expression of the electron transport chain, oxidative phosphorylation machinery, and fatty acid oxidation regulation, we performed Western blot analysis in cells treated with red light at 80% confluence. Cells were lysed over ice in the presence of RIPA buffer containing protease and phosphatase inhibitor cocktails, and total protein levels were quantified using the Pierce BCA Protein Assay kit. Lysates were prepared in sample buffer (2% SDS, 10% glycerol, 0.062 M Tris pH 6.8, 0.002% bromophenol blue, 5% 2-mercaptoethanol), 40 ug of total protein were loaded onto 10% SDS-PAGE, and electro-transferred to PVDF membranes. Membranes were blocked with 5% BSA in TTBS (20 mM Tris pH 7.5, 150 mM NaCl, and 0.1% Tween 20) for 1 h at room temperature before incubation with Total OXPHOS Human WB Antibody Cocktail (Abcam:110411, 1:1000), CPT1 (Santa Cruz: 393070, 1:1000), CPT2 (Abcam: 181114, 1:500), AMPK (Cell signaling: 5831S, 1:1000), phospho-AMPK (Abcam: 133448, 1:1000), PDH (Abcam: 110330, 1:1000), ACC (Cell Signaling: 3676, 1:1000), phospho-ACC (Cell Signaling: 11818, 1:500), ETFA (HPA018990, 1:1000); primary antibodies where incubated at 4°C overnight. Fluorescent 660 nm secondary anti-mouse or anti-rabbit antibodies (1:20.000) were incubated for 1 h at room temperature prior to fluorescence detection using a ChemiDoc Imaging System (Bio-Rad). Quantification of band densitometry was performed using the ImageLab software. Given that no single protein is a strong housekeeping control in metabolism, protein expression values obtained were normalized per pixels of total protein in each well, as evidenced by Ponceau S dye (Sigma).

### 2.8. Lipid Measurements

Cells were seeded and incubated until 80% confluence on six-well plates in 2 mL of high glucose DMEM with 10% FBS, 1% Penicillin/Streptomycin, without phenol red. Cells were treated with different doses of red light (0, 12 and 36 J/cm^2^) and 2 hours post treatment cells were PBS-washed twice and scraped in 1 mL PBS; from this solution 800 μL were centrifuged and 200 μL were kept for normalization by protein quantification. After centrifugation in 5 min per 300 g, we extracted the lipids following an adapted Folch’s method [15, 16]. Briefly, cell pellets were suspended in 1 mL chloroform:methanol (2:1) and vigorously agitated for 1 hour at room temperature. 200 μL of MilliQ water were added to the tube and vortexed to separate the homogenate in two phases. We collected the organic phase into fresh microcentrifuge tubes and evaporated the solvent overnight in a fume hood overnight. The dried lipid extract was then suspended in 200 μL of chloroform containing 2% Triton X-100 and allowed it to dry completely at room temperature for 2 hours (or until complete solvent evaporation). The resulting lipid film was suspended in 50 μL of Milli-Q water and quantified using the following commercial assay kits according to manufacturers protocols: Triglycerides (LabTest®, Brazil), Cholesterol (LabTest®, Brazil), and Non-Esterified Fatty Acids (Wako Diagnostics, Fujifilm, USA). All measured values were normalized to total cellular protein content.

### 2.9. Statistical Analysis

Data are presented as individual biological replicates with mean ± standard deviation (SD) from at least three independent experiments. All statistical analyses were performed using GraphPad Prism 8 (GraphPad Software, San Diego, CA). For high-resolution respirometry comparisons between two conditions, we applied two-tailed Student’s t-tests with Welch’s correction for unequal variances. Multiple comparisons involving one independent variable were analyzed by one-way ANOVA followed by Tukey’s post-hoc or Dunnett’s tests. Statistical significance was defined as p < 0.05.

## 3. Results

### 3.1. Effect of light on keratinocyte viability is dependent on wavelength

To initially assess the effect of UVA and visible light on keratinocytes, we selected the midpoints of the spectra (i.e., 365 nm, 450 nm, 517 nm, and 660 nm, see Supplementary Figure S1 for spectra) and submitted cells to one dose of 36 J/cm² at each wavelength. Viability was measured using MTT, Cristal Violet staining (CVS), and Neutral Red staining (NRS), so as not to incur on specific light effects on the specific processes assessed by each of these dyes (Fig. 1). UVA exposure resulted in loss of cell viability 48 hours after exposure as measured by all methods (Fig. 1A-C), an effect maintained 72 hours post treatment (Fig. 1D-F) and compatible with prior findings [15]. Blue light had a noticeable effect at 24 hours compared to non-irradiated controls, but without statically significance. Interestingly, blue and green light promoted a decrease in MTT and CVS at 48 hours (Fig. 1A-B) and 72 hours (Fig. 1D, E), which indicates that 36 J/cm^2^ of green light inhibits keratinocyte growth. Conversely, red light-treated cells displayed enhanced CVS intensity, suggesting it increased proliferation at 48 hours (Fig. 1B) and 72 hours post treatment (Fig. 1E). Overall, the results in Fig. 1 indicate that keratinocytes respond differently to the same dose of light at distinct wavelengths, with responses ranging from overt cell death (UVA light) to augmented cell counts (red light).

**Figure 1.**
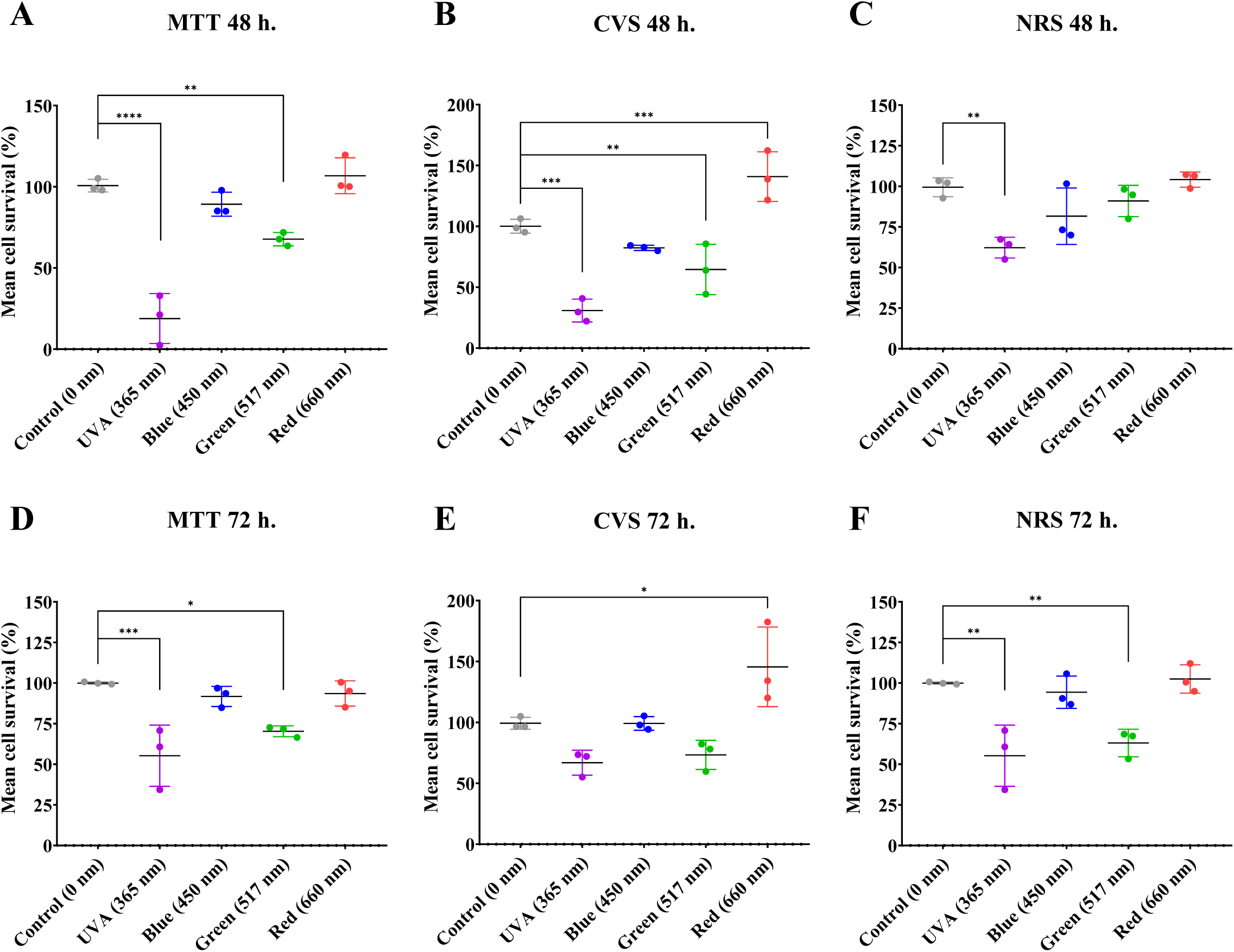
The effect of light on cell viability in keratinocytes is dependent of wavelength. Cell viability was quantified after 48 and 72 hours post-irradiation at 36 J/cm^2^ with UVA (365 nm), blue light (450 nm), green light (517 nm), or red light (660 nm). Viability was assessed using: MTT, Cristal Violet Staining (CVS), or neutral red staining (NRS). Results are expressed as means ± SD of three independent experiments; ns = > 0.1. *p = <0.0332. **p = 0.0021. ***p = 0.0002 ****p = <0.0001, One-way ANOVA followed by Dunnett.

### 3.2. Metabolic fluxes are affected by different light wavelengths

To investigate the implications of light on electron transport and oxidative phosphorylation, an expected target of irradiation due to their dependence on various light absorbing molecules, we measured metabolic fluxes through oxygen consumption rates (OCRs) in intact cells exposed to 36 J/cm^2^ of different wavelengths of light. Intending to uncover metabolic changes induced early on in cells, OCRs were measured 2 hours post irradiation in plated cells using a Seahorse Extracellular Flux Analyzer (Fig. 2 and Fig. S2). Figure 2A shows typical intact cell OCR measurements over time. Measurements were made under basal conditions (prior to any additions) and reflecting the normal metabolic status of the cells (quantified in Fig. 1B). Oligomycin, an ATP synthase inhibitor, was then added to quantify OCRs dependent on mitochondrial oxidative phosphorylation (quantified in Fig. 1C). Maximal electron transport was then induced by adding the uncoupler CCCP (quantified in Fig. 1D), followed by antimycin and rotenone to inhibit mitochondrial electron transport and determine nonmitochondrial OCRs, which were subtracted from all quantifications.

**Figure 2.**
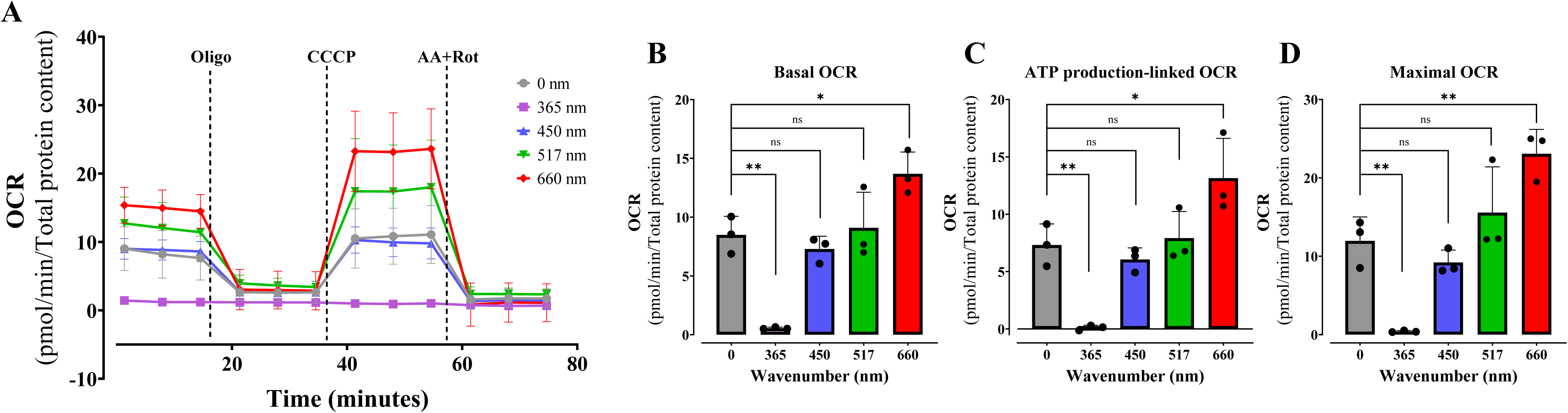
Mitochondrial respiration is differentially modulated by different light wavelengths. Cell oxygen consumption rates (OCR) were quantified after 2 hours irradiation at 36 J/cm^2^ with UVA (365 nm), blue light (450 nm), green light (517 nm), or red light (660 nm) in keratinocytes. OCRs were measured as described in Methods under basal conditions, followed by injection of oligomycin (oligo, 1 μM), CCCP (1 μM), and antimycin A plus rotenone (AA/Rot, 1 μM each). Results are expressed as means ± SD of three independent experiments; ns = > 0.1. *p = <0.0332. **p = 0.0021, One-way ANOVA followed by Tukey.

Compared to controls (gray lines/bars), we found that UVA irradiation almost completely abrogates OCRs (Fig. 2A-D), a result compatible with the loss of viability seen in Fig. 1. Blue and green light promoted no changes in basal (Fig. 2B), ATP production-linked (Fig. 2C), or maximal respiration (Fig. 2D). Interestingly, keratinocytes presented a substantial increase in all OCR states when treated with red light: basal respiration (Fig. 2B), ATP production (Fig. 2C), and maximal respiration (Fig. 2D) were enhanced, demonstrating that red light significantly increases oxidative phosphorylation and electron transport capacity. Extracellular acidification rates (ECARs) and OCRs associated to the proton leak, reserve capacity and non-mitochondrial respiration were equal between all treatments, except for enhancements promoted by red light (Supplementary Fig. S2). Overall, we find that metabolic fluxes in intact cells are impacted by light differentially depending on wavelengths, with a highly interesting enhanced basal, ATP-linked, and maximal oxygen consumption observed after red light irradiation.

### 3.3. Red light increases oxidative phosphorylation within a physiological dose range, with long-lasting effects

We then sought to better characterize the enhancement of oxidative phosphorylation seen with red light, and tested if different doses of 660 nm irradiation had similar effects (Fig. 3 and Fig. S3). We found that increasing exposure up to 36 J/cm^2^ enhanced OCRs dose-dependently, but that a higher exposure of 150 J/cm^2^ decreased OCRs, probably due to photothermal or phototoxic effects. Importantly, red light doses between 5 and 150 J/cm^2^ are compatible with typical human exposure and with photobiomodulation protocols [7, 16], indicating that the modulatory effects seen here could constitute a physiological response in vivo.

**Figure 3.**
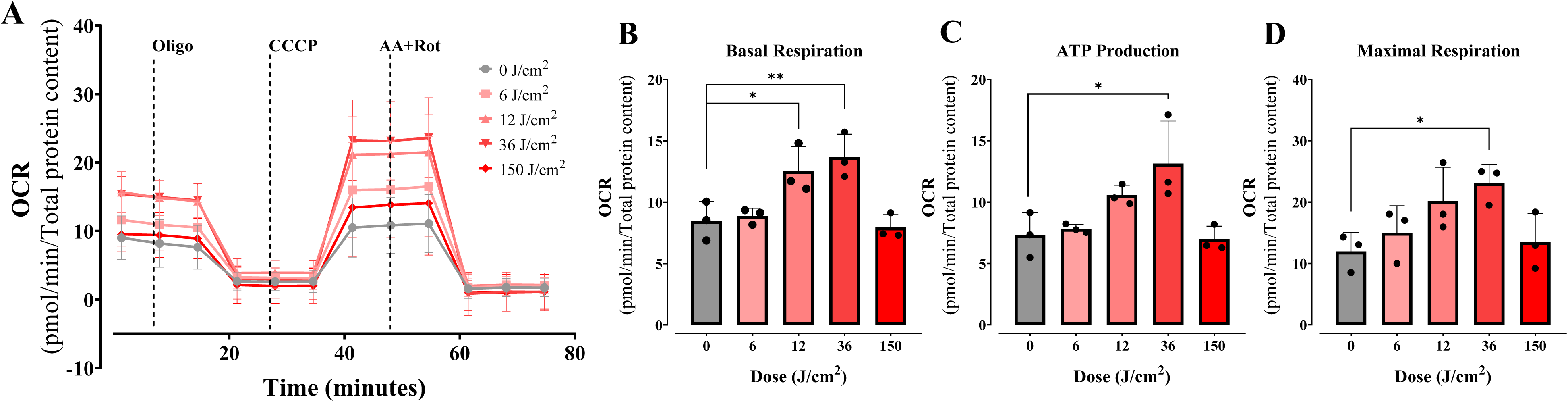
Moderate doses of red light (660 nm) increase mitochondrial respiration. OCRs were quantified under similar conditions to Fig. 2 after 2 hours irradiation at 6, 12, 36 and 150 J/cm^2^ with red light in keratinocytes. ns = > 0.1. *p = <0.0332. **p = 0.0021. ***p = 0.0002. ****p = <0.0001. Results are expressed as mean ± SD of three independent experiments; One-way ANOVA followed by Dunnett.

Although all our experiments were conducted in temperature-controlled conditions, the irradiator’s refrigeration system may be less able to maintain local temperatures than ambient chamber temperatures. We thus measured plate heating using infrared thermography, which revealed some heating at higher red light doses (Fig. 4A: 12-36 J/cm² produced physiological temperature ranges (mean 37°C ± 0.5°C, max 38°C ± 0.5°C), whereas high irradiation doses (150 J/cm²) reached 40°C, which could in theory explain the inhibitory effects observed in our initial assays (Fig. 3). To isolate thermal contributions, we conducted dark-control experiments at 37 and 38°C (matching effective photobiomodulation temperatures) using identical OCR protocols. No significant changes occurred in basal, ATP-linked or maximal respiration occurred (Fig. 4B-E).

**Figure 4.**
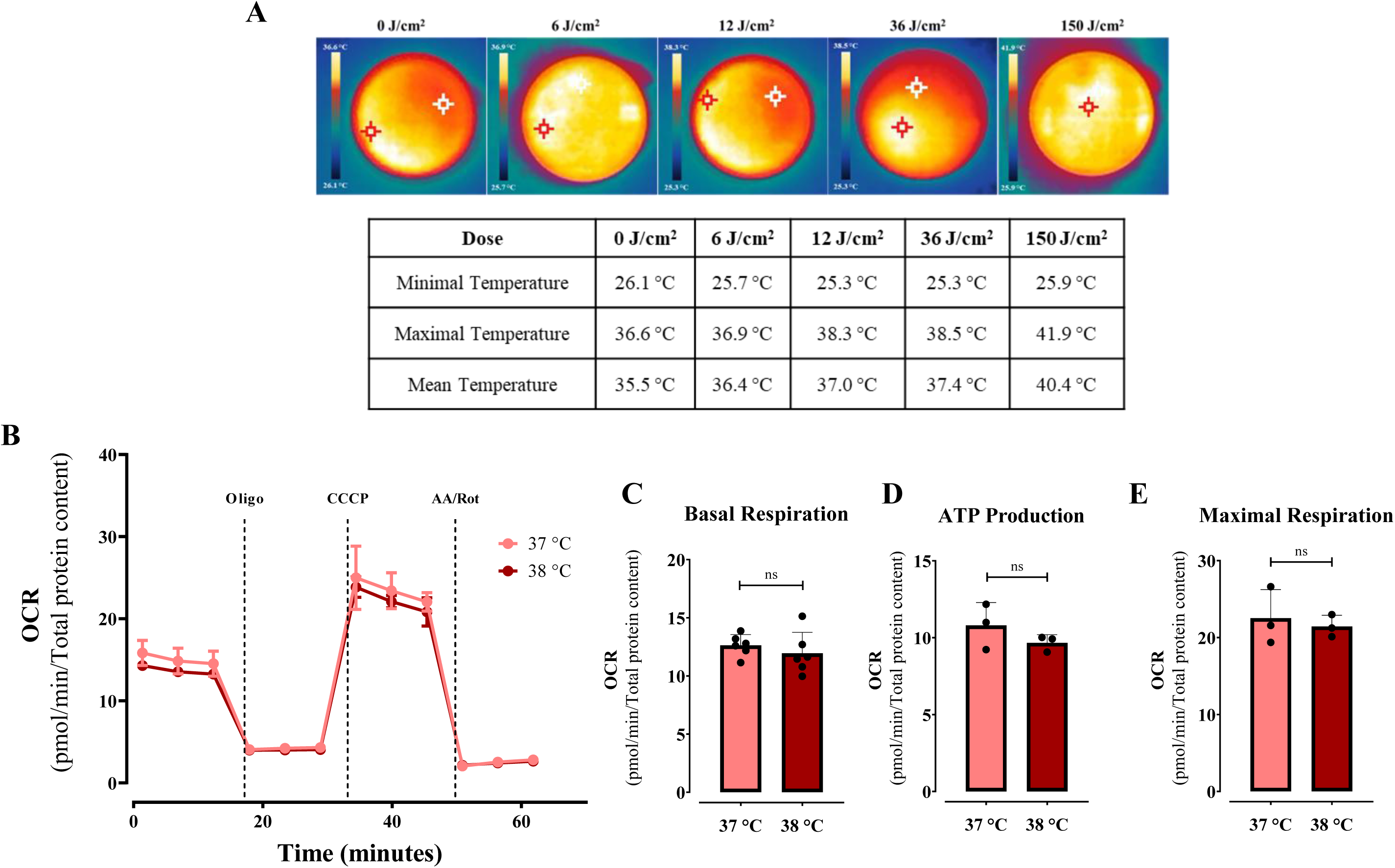
Red light effect is not photothermal. Local temperature measurements were detected using an infrared camera. Minimal, mean and maximal temperature were calculated automatically by pixel intensity in each dose of light.

As we had measured a single time point (two hours after irradiation) so far, we sought next to uncover the timeline of the enhanced oxygen consumption rates observed in our cells. For this, we used a Resipher Real-time Cell Analyzer (Lucid Scientific, Atlanta, GA, USA), capable of measuring oxygen consumption in plated cells within normal growth conditions in normal laboratory CO_2_ incubators. Cells were plated under conditions similar to those used in the Seahorse metabolic flux experiments, except that OCRs were quantified continuously immediately after a single bought of red-light exposure at different intensities (Fig. 5). We found that oxygen consumption rates became elevated soon after irradiation, in a dose-dependent manner, a result compatible with the Seahorse data, although higher, 150 J/cm^2^ doses also elevated OCRs in this setting, possibly due to the presence of bicarbonate (the only discernable experimental difference between experimental setups). Surprisingly, this effect was long-lasting, and maintained for over 2 days, irrespective of daily changes in culture media, which were accompanied by a temporarily decreased OCR. Starting at day 3 post-irradiation, OCRs were again similar in irradiated and non-treated groups. The finding that red light enhancement of oxygen consumption rates and capacity is long-lasting is interesting, and suggests mechanisms other than direct light interactions.

**Figure 5.**
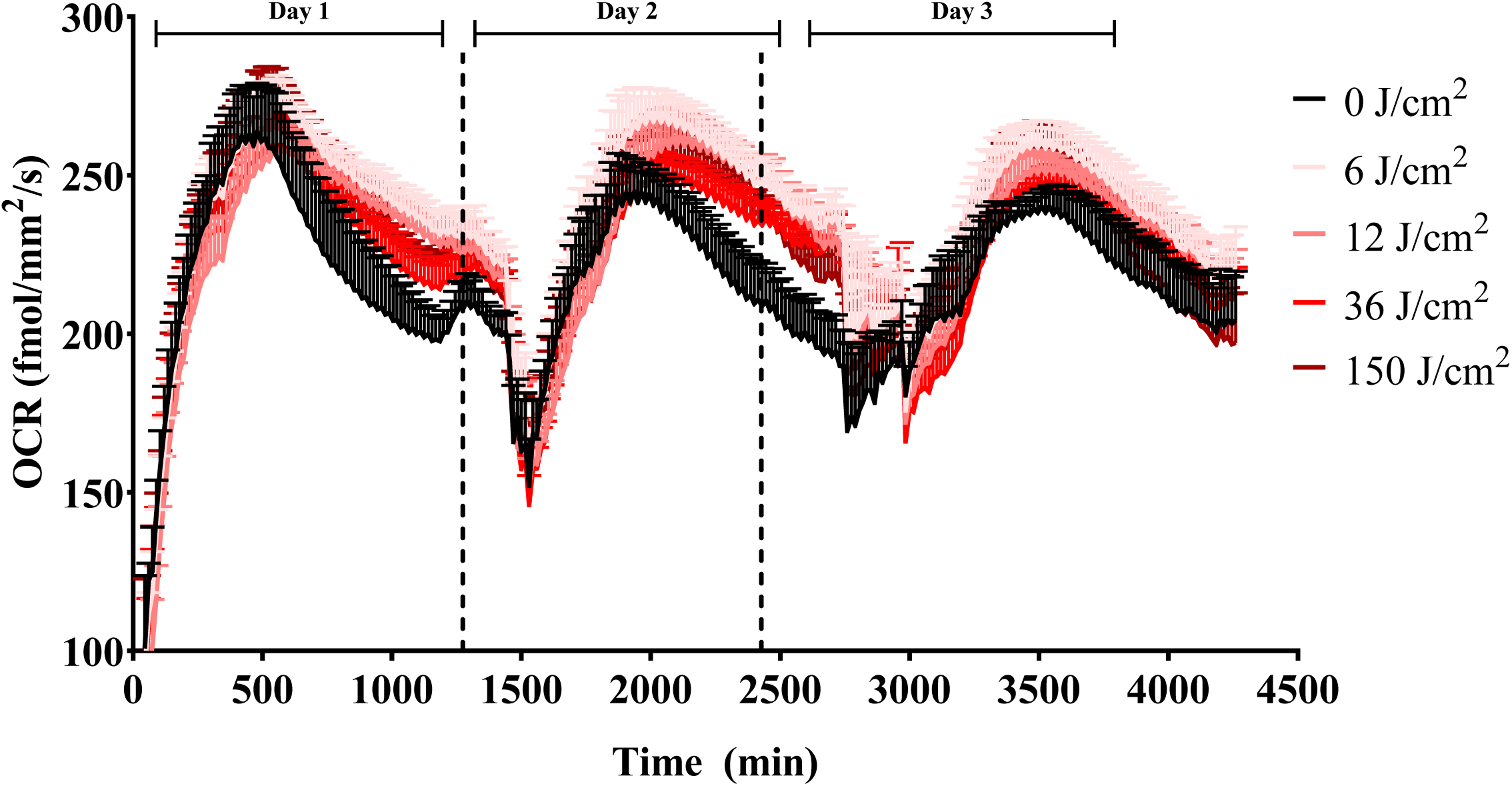
Increased oxygen consumption promoted by red light (660 nm) is prologued. OCRs were measured over three days in culture using a Resipher Real-Time Cell Analyzer under basal conditions. Keratinocyte respiration (OCR) quantification was initiated immediately post irradiation with red light at different doses (0, 6, 12, 36 and 150 J/cm^2^). Results are means + SEM of 4 biological replicates.

Although our study has primarily focused on keratinocytes, we extended our investigation to fibroblasts (Hs68) and melanocytes (B16-F10), to determine cell specificity of red light effects, performing identical Seahorse XF Mito Stress Tests, 2 hours after red light irradiation at varying doses. Strikingly, fibroblasts showed no significant OCR changes at any dose (Fig. 6A-D, S4), while melanocytes exhibited dose-sensitive responses, with low doses enhancing basal respiration (Fig. 6F) and ATP-linked OCR (Fig. 6G) but not maximal respiration (Fig. 6H). These results contrast with keratinocytes, where effects were observed at moderate doses.

**Figure 6.**
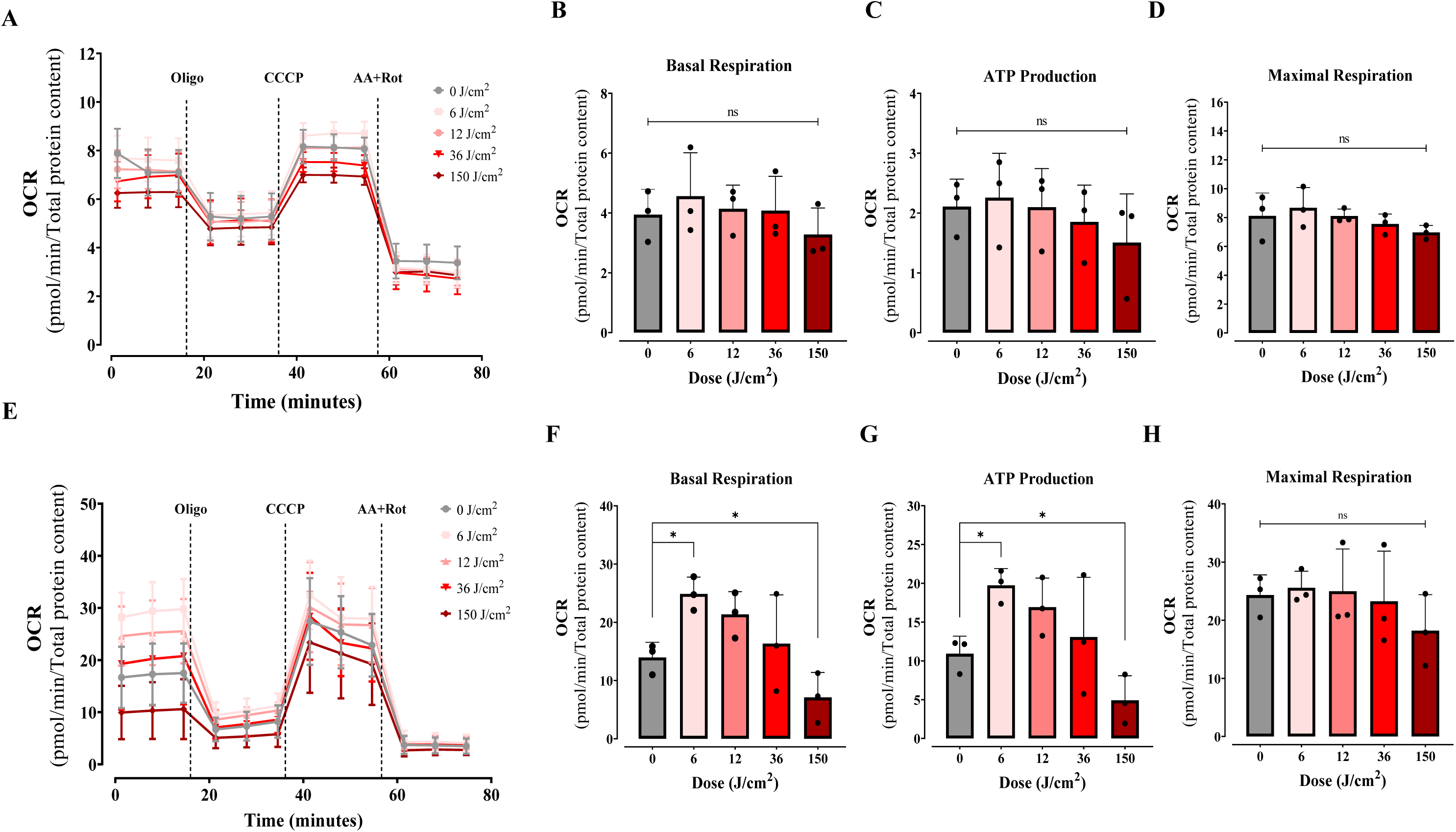
Red light stimulation of OCRs is cell-specific. Fibroblasts (A-D) and melanocytes (E-H) were treated under similar conditions to Fig. 2 after 2 hours of irradiation at 6, 12, 36, and 150 J/cm^2^ with red light (660 nm) and OCRs were quantified. Results are expressed as means ± SD of three independent experiments; p < 0.05, one- way ANOVA followed by Dunnett’s test.

### 3.4. Red light does not alter mitochondrial oxidative phosphorylation machinery

We sought next to identify the mechanisms in which red light promotes long-lasting enhanced OCRs in keratinocytes. We began by analyzing the quantities of respiratory protein complexes (OXPHOS complex subunits) by western blots (Fig. 7), and observed small decreases in Complex I – NADH dehydrogenase (I-NDUFBS), but no significant differences in the Complex II – Succinate dehydrogenase (SDHB), Complex III - Q-cytochrome c oxidoreductase (III-UQCRC), Complex IV – cytochrome C oxidase (IV- COX II) or Complex V - ATP Synthase (VATP5A), relative to the dark control.

**Figure 7.**
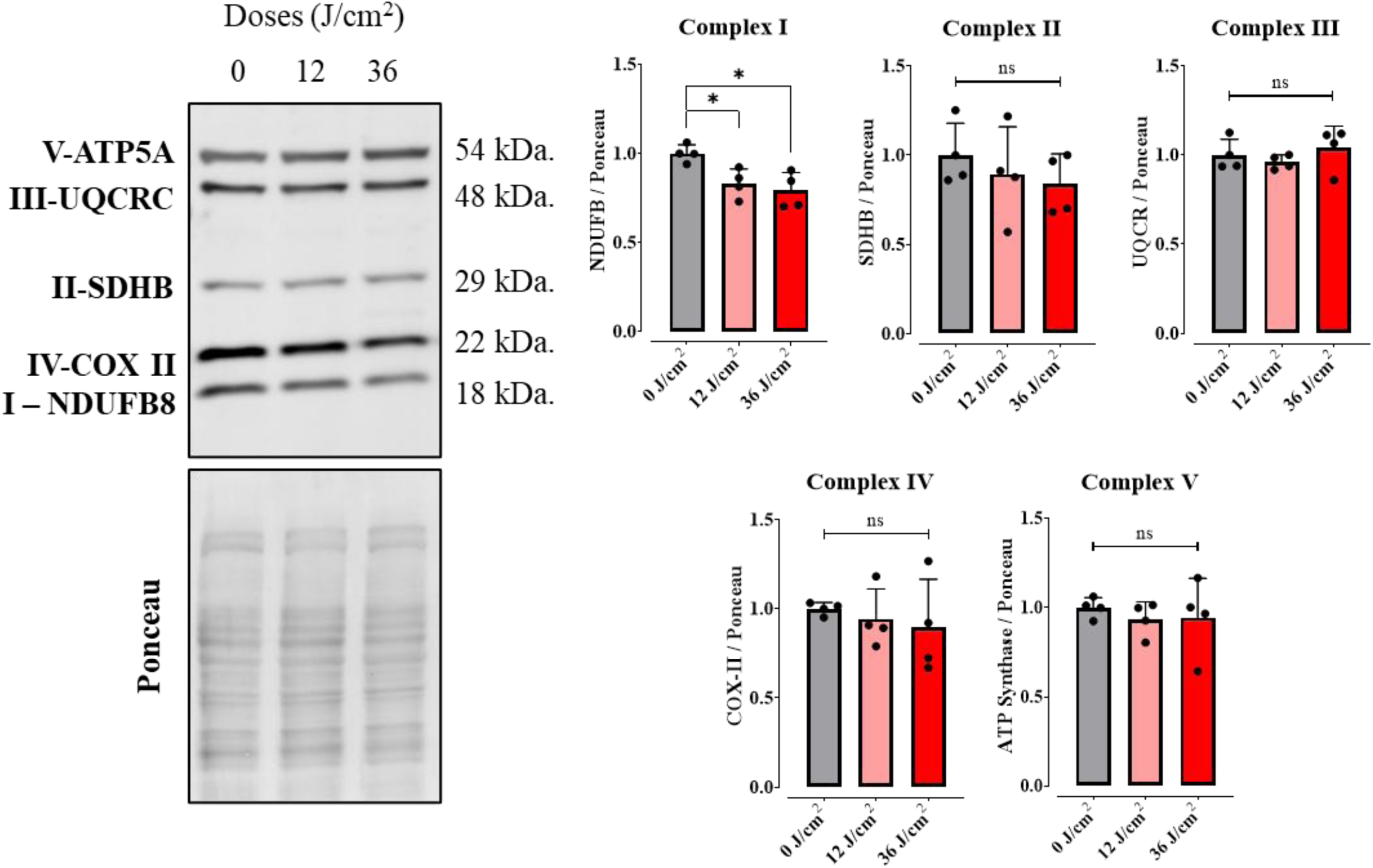
Red light does not increase mitochondrial respiratory complex expression. Western blot analysis of electron transport chain and oxidative phosphorylation protein levels was conducted in cells 2 hours post exposure to red light at different doses (0, 12, 36 J/cm^2^). Extracts were collected and protein quantified by BCA. Complex I – NADH dehydrogenase (I-NDUFBS), Complex II – Succinate dehydrogenase (SDHB), Complex III - Q-cytochrome c oxidoreductase (III-UQCRC), Complex IV – Cytochrome C oxidase (IV-COX II) and Complex V - ATP Synthase (V-ATP5A). ns = > 0.1. *p = <0.0332. Results are expressed as means ± SD of four independent experiments; One-way ANOVA followed by Tukey.

The activity of oxidative phosphorylation components may be altered independently of their expression. Indeed, some groups have suggested that red light directly activates cytochrome c oxidase [17, 18]. While our finding that the effects of red light are long-lasting is not compatible with this mechanism (which would require the presence of light during measured activation), we decided to measure electron transport rates more directly in permeabilized cells (Fig. 8), in which cytoplasmic factors are infinitely diluted, allowing us to directly probe mitochondrial oxygen consumption regulation using a Clark-type high resolution respirometer (Oroboros Oxygraph-2k). Oxygen consumption supported by a physiological mixture of substrates was measured under initial basal conditions and after the addition of ADP (which stimulated oxidative phosphorylation), oligomycin (which inhibits oxidative phosphorylation), and CCCP (which promotes uncoupling and maximizes electron transport). Respiratory control ratios (ADP-induced/oligomycin-treated OCRs) were also calculated. None of these parameters were significantly altered by red light. This clearly indicates that the respiratory enhancement observed in intact cells is not centered on a mitochondrial regulatory mechanism, and therefore probably involves modulation of cytosolic metabolic pathways upstream of oxidative phosphorylation.

**Figure 8.**
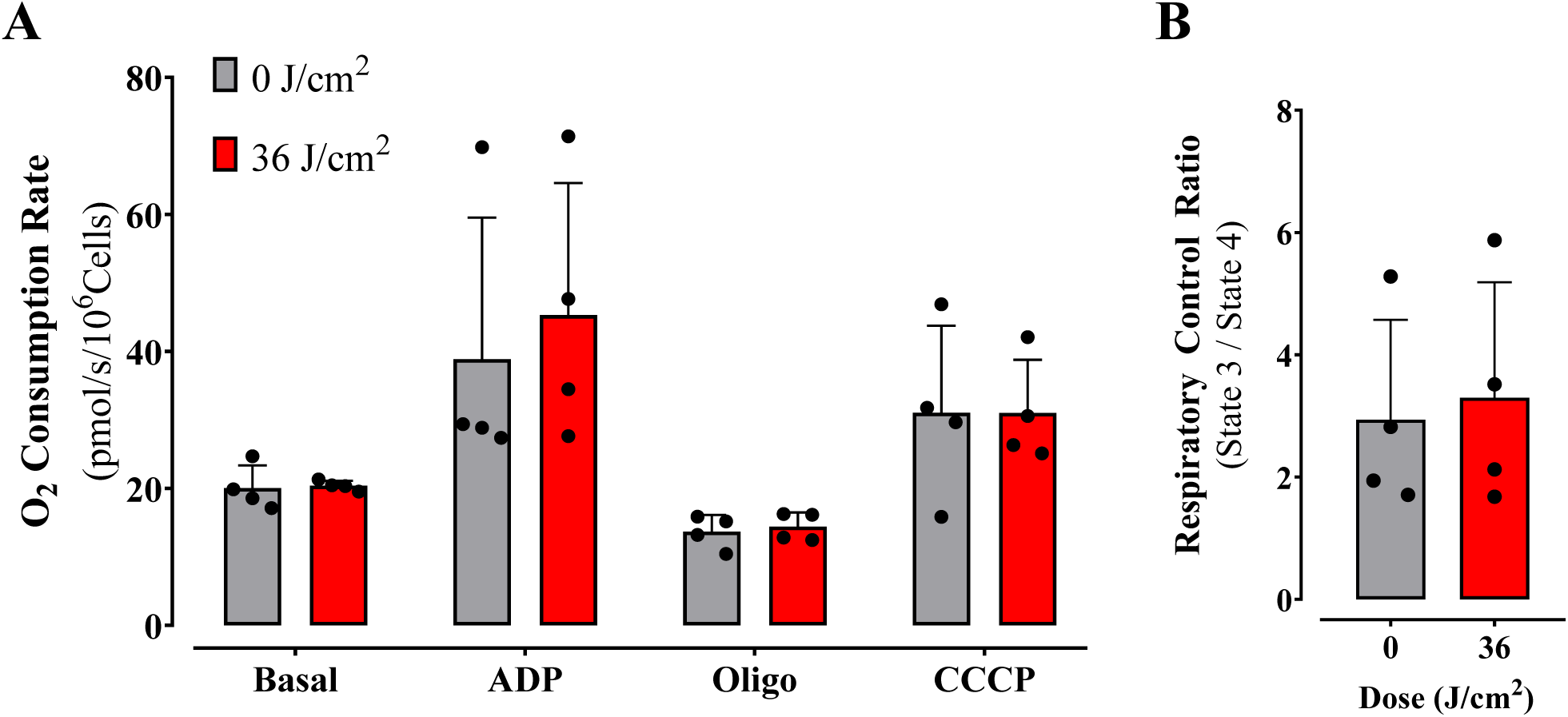
Effect of red light on mitochondrial oxidative phosphorylation requires intact cells. A) Permeabilized cells were incubated in the presence of 0.0017% digitonin, 1 mM succinate, 1 mM pyruvate, and 1 mM malate (state 2), 1 mM ADP (state 3), 1 μM oligomycin (Oligo, state 4), and 1 mM CCCP (state 3U), added successively while oxygen consumption was monitored as described in Methods. B) respiratory control ratios were calculated as the ratio between state 3 and state 4. Results are means + SD of 3 biological replicates.

### 3.5. Red light irradiation specifically enhances fatty acid oxidation in keratinocytes

Since mitochondrial OCRs were equal in irradiated and control cells when substrates were delivered directly to the organelles of permeabilized cells, but enhanced by light in intact cells, we investigated if cytosolic pathways fueling mitochondrial oxidative phosphorylation were affected by red light.

Glucose metabolism was dynamically assessed first, by quantifying proton release dependent on glucose degradation and enhanced by oligomycin, an ATP synthase inhibitor which promotes ATP depletion and activates the pathway (Fig. 9). By following ECARs over time after the additions of glucose, oligomycin and the competitive inhibitor 2-deoxiglucose (Fig. 9A), we were able to quantify basal (Fig. 9B), maximal (Fig. 9C), and reserve (Fig. 9D) glycolytic rates in irradiated and control cells, as well as non-glycolytic acidification rates (Fig. 9E). Results clearly show that there is no effect of red light, as none of these parameters are significantly altered.

**Figure 9.**
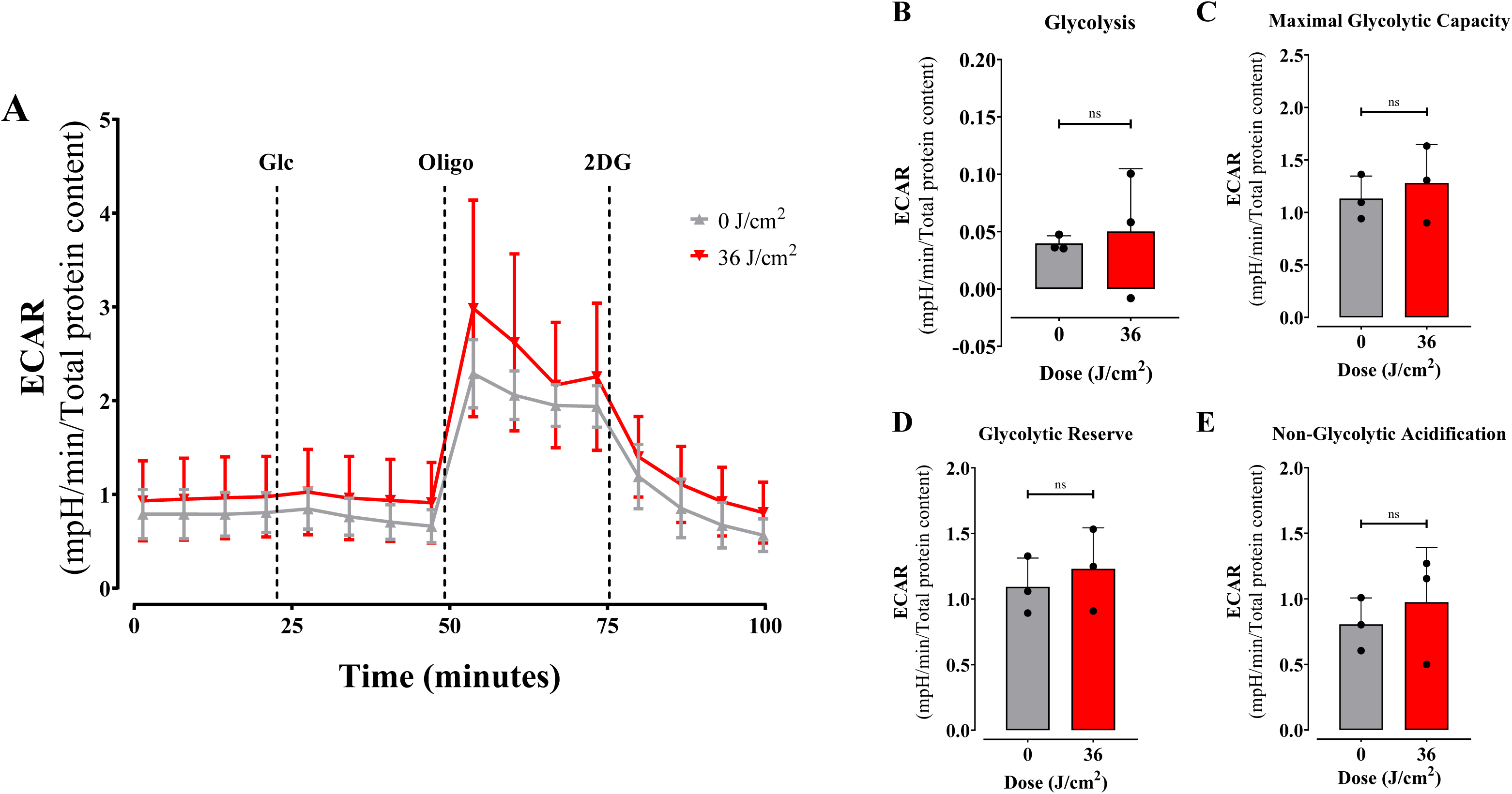
Glycolysis is not affected by red light. Extra Cellular Acidification Rates (ECARs) were measured as described in Methods in intact cells 2 hours post irradiation at 36 J/cm^2^ with red light (660 nm) under basal conditions, followed by injection of Glucose (Glc, 25 mM), Oligomycin (Oligo, 1 μM) and 2-Deoxy-Glucose (2DG, 50 mM). ns = > 0.1. Results are expressed as means ± SD of three independent experiments; Student’s t test.

Next, we measured the effect of red light on amino acid metabolism, by following the effect of BPTES, a glutaminase inhibitor, on OCRs in the presence and absence of red light. BPTES was added to control or red light-irradiated keratinocytes, and OCRs were followed over extended time to allow for glutaminase inhibition (Fig. 10A). Basal, maximal, and ATP linked respiration were enhanced by red light, but this effect was not abrogated by BPTES, indicating that it does not involve a change in glutaminase activity (Fig. 10B-E).

**Figure 10.**
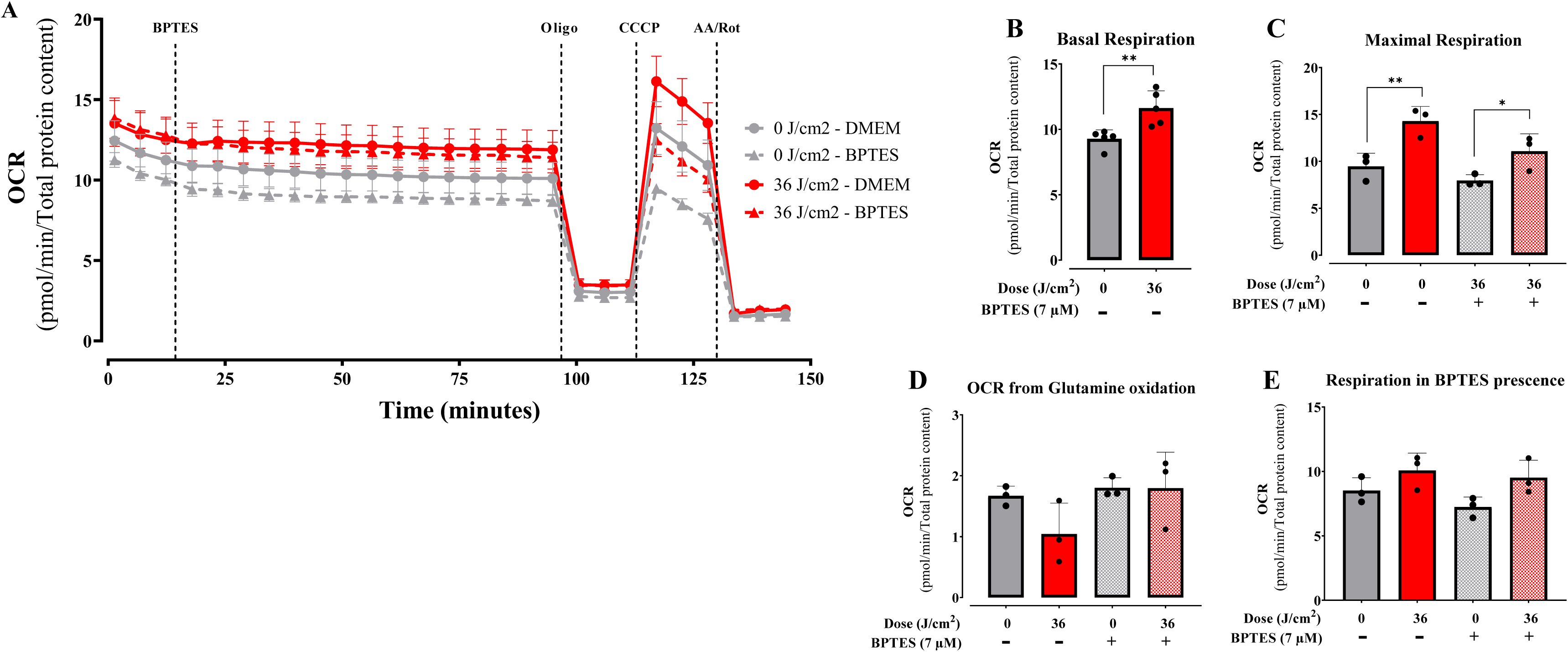
Glutamine oxidation is unaffected by red light. Oxygen Consumption Rates sensitive to glutaminase inhibitor BPTES was analyzed 2 hours post irradiation at 36 J/cm^2^ with 660 nm under basal conditions, followed by injection of BPTES (7 μM) oligomycin (oligo, 1 μM), CCCP (1 μM), and antimycin A / rotenone (AA/Rot, 1 μM each). Results are expressed as means ± SD of three independent experiments; *p = 0.0021, Student’s t test. ns = > 0.1. *p = <0.0332 One-way ANOVA followed by Tukey.

Next, we measured oxygen consumption promoted by fatty acid oxidation (FAO), specifically long-chain fatty acids, which are the main lipid energy source in mammalian cells. To measure FAO-dependent metabolic fluxes, we compared OCRs before and after the addition of etomoxir, an inhibitor of CPT1, which transports fatty acids into mitochondria, where they are degraded by fatty acid oxidation. Similarly to CCCP and oligomycin, the use of etomoxir was preceded by careful calibration experiments to determine ideal quantities and avoid off-target effects such as uncoupling (which increases OCRs in the presence of oligomycin). Using calibrated etomoxir quantities, we found that OCRs were decreased by etomoxir over time specifically in red light-irradiated cells (Fig. 11), while inhibition of FAO had a non-significant effect on control cells. Indeed, etomoxir abrogated the stimulated basal (Fig. 11B) and maximal (Fig. 11C) OCRs observed in red light-irradiated cells, indicating that increased CPT1- dependent FAO is the mechanism in which red light promotes enhanced oxygen consumption in intact cells.

**Figure 11.**
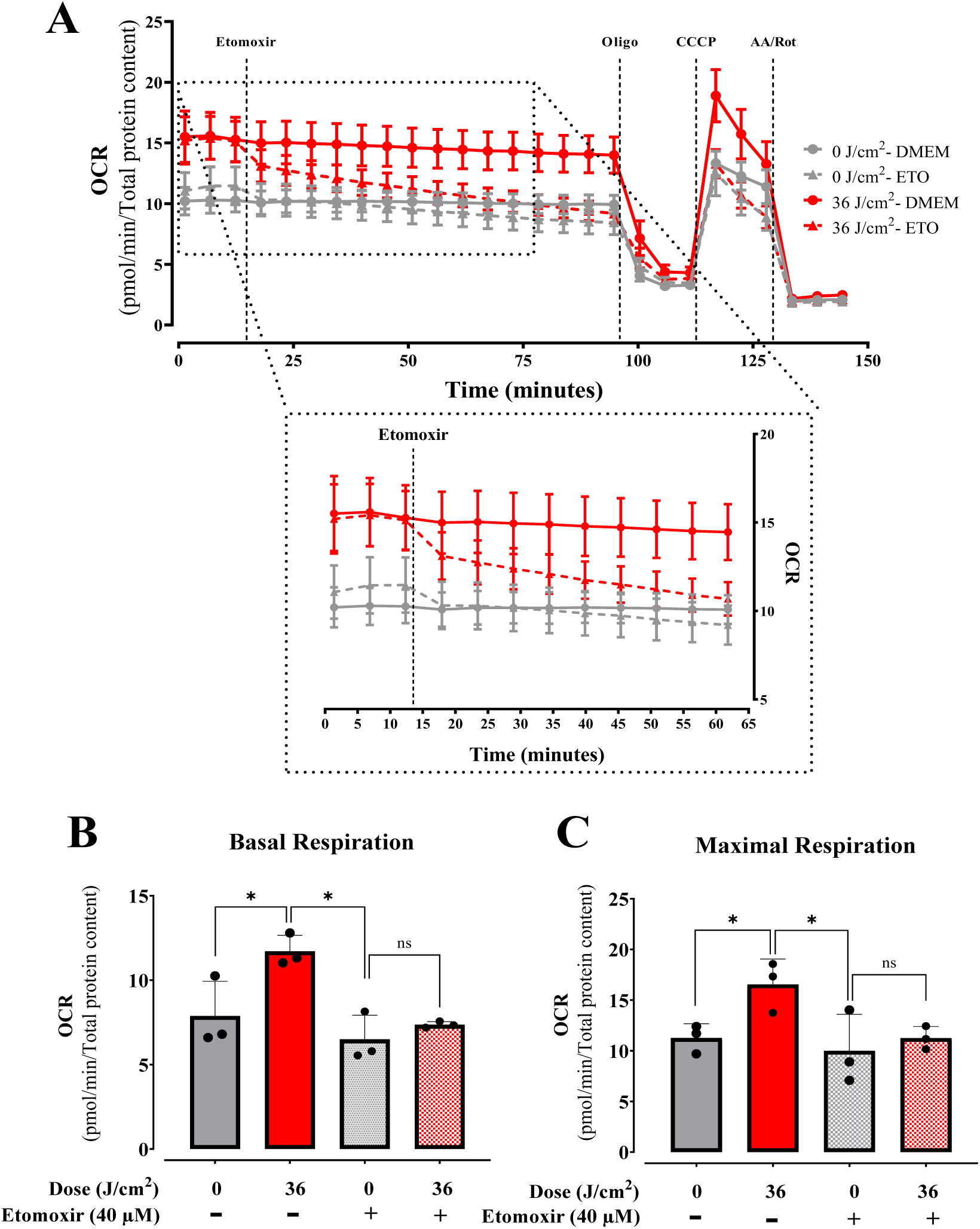
Fatty acid oxidation is enhanced by red light. Oxygen Consumption Rates sensitive to CPT1 inhibitor etomoxir were analyzed 2 hours post irradiation at 36 J/cm^2^ with 660 nm under basal conditions, followed by injection of Etomoxir (40 μM), oligomycin (oligo, 1 μM), CCCP (1 μM), and antimycin A / rotenone (AA/Rot, 1 μM each). Results are expressed as means ± SD of three independent experiments; *p = 0.0021, Student’s t test. ns = > 0.1. *p = <0.0332 One-way ANOVA followed by Tukey.

### 3.6. Red light decreases cellular free fatty acid content by activation AMPK/ACC phosphorylation

Given our finding that red light specifically increased fatty acid oxidation-dependent OCRs, we quantified lipids in our cells. We found that that red light specifically reduced free fatty acid (FFA) levels within 2 hours of irradiation (Fig. 12A), while triglyceride (Fig. 12B) and cholesterol (Fig. 12C) concentrations remained unchanged. This selective FFA reduction, coupled with the etomoxir-sensitive OCR enhancement, strongly supports red light’s ability to stimulate β-oxidation. To determine if this metabolic shift involved upregulation of mitochondrial fatty acid utilization machinery, we analyzed key proteins including fatty acid transporters (CPT1, CPT2), acetyl-CoA-generating enzyme PDH, and electron transfer component ETFA (Fig. S4). None showed altered expression relative to dark controls. These results demonstrate that red light’s metabolic effects occur without remodeling mitochondrial protein content, instead implicating acute cytosolic regulation as the primary control mechanism for enhanced fatty acid oxidation.

**Figure 12.**
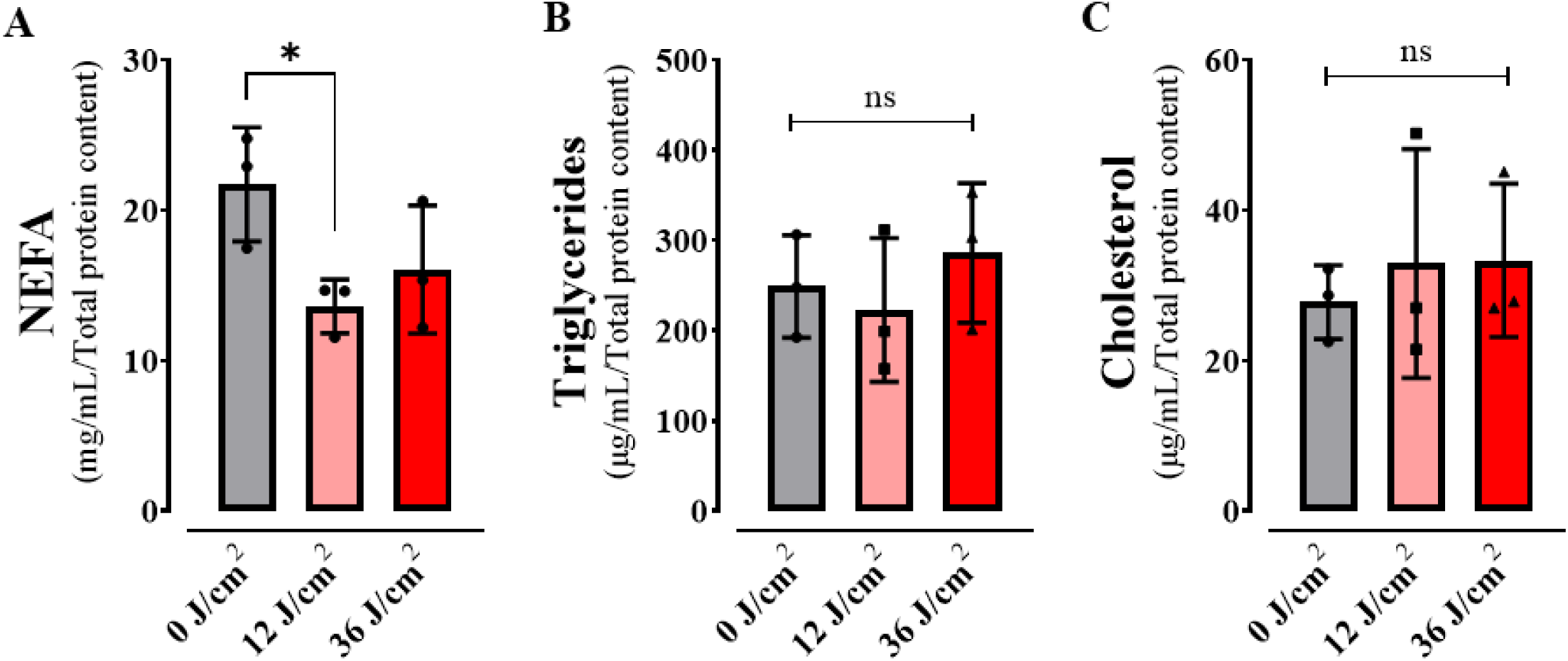
Red light reduces free fatty acid content. Non-Esterified Free Fatty Acids (A), Triglycerides (B) and Cholesterol (C) where quantified in keratinocytes after 2 hours post red light in doses of 0, 12 ad 36 J/cm^2.^ Results are expressed as means ± SD of three independent experiments; ns = > 0.1. *p = <0.0332, One-way ANOVA followed by Dunnett.

To test our hypothesis of cytosolic metabolic regulation, we investigated two key controllers of fatty acid oxidation (FAO): AMP-activated protein kinase (AMPK) and its downstream target acetyl-CoA carboxylase (ACC). Western blot analysis revealed that while red light did not alter total AMPK (Fig. 13A) or ACC (Fig. 13D) protein levels, it significantly increased phosphorylation of both enzymes (P-AMPK, Fig. 13B; P-ACC, Fig. 13E), elevating P-AMPK/AMPK (Fig. 13C) and P-ACC/ACC (Fig. 13F) ratios. These findings demonstrate that red light activates FAO through post-translational modification rather than protein abundance changes. Mechanistically, AMPK phosphorylation likely induces ACC inhibition, simultaneously blocking fatty acid synthesis while promoting free fatty acid (FFA) flux toward mitochondrial β-oxidation.

**Figure 13.**
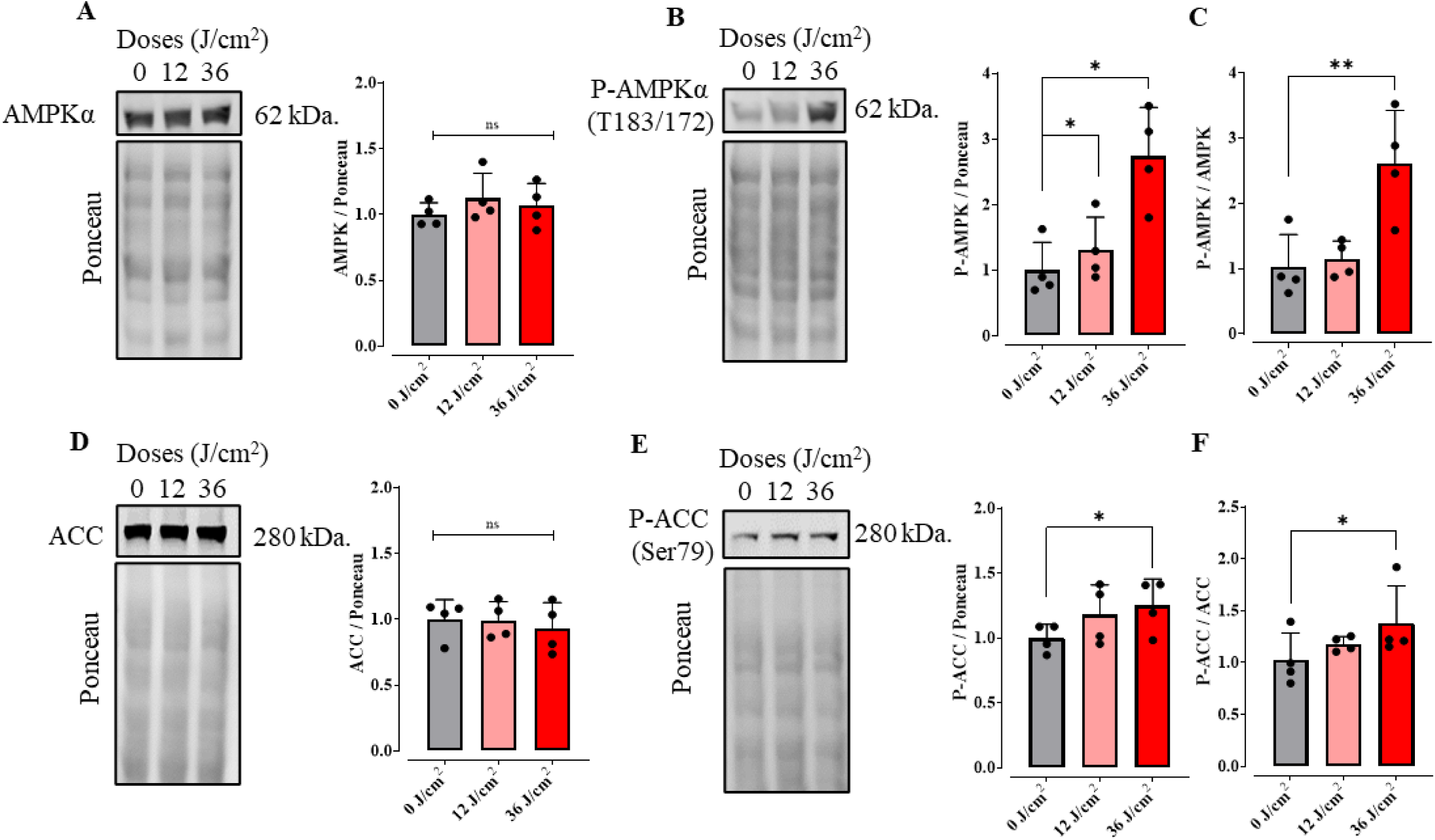
Red light activates Fatty acid regulatory steps. Western blot analysis of protein levels was conducted in cells 2 hours post exposure to red light at different doses (0, 12, 36 J/cm^2^). Extracts were collected and protein quantified by BCA. AMP-activated protein kinase (AMPK) (A), Acetyl-CoA carboxylase (ACC) (D), and their phosphorylated forms where quantified (B, E) and the ratio of P-AMPK/AMPK (C) and P-ACC/ACC (F) were calculated. Results are expressed as means ± SD of four independent experiments; ns = > 0.1. ns = > 0.1. *p = <0.0332. **p = 0.0021. One-way ANOVA followed by Dunnett.

**Figure 14.**
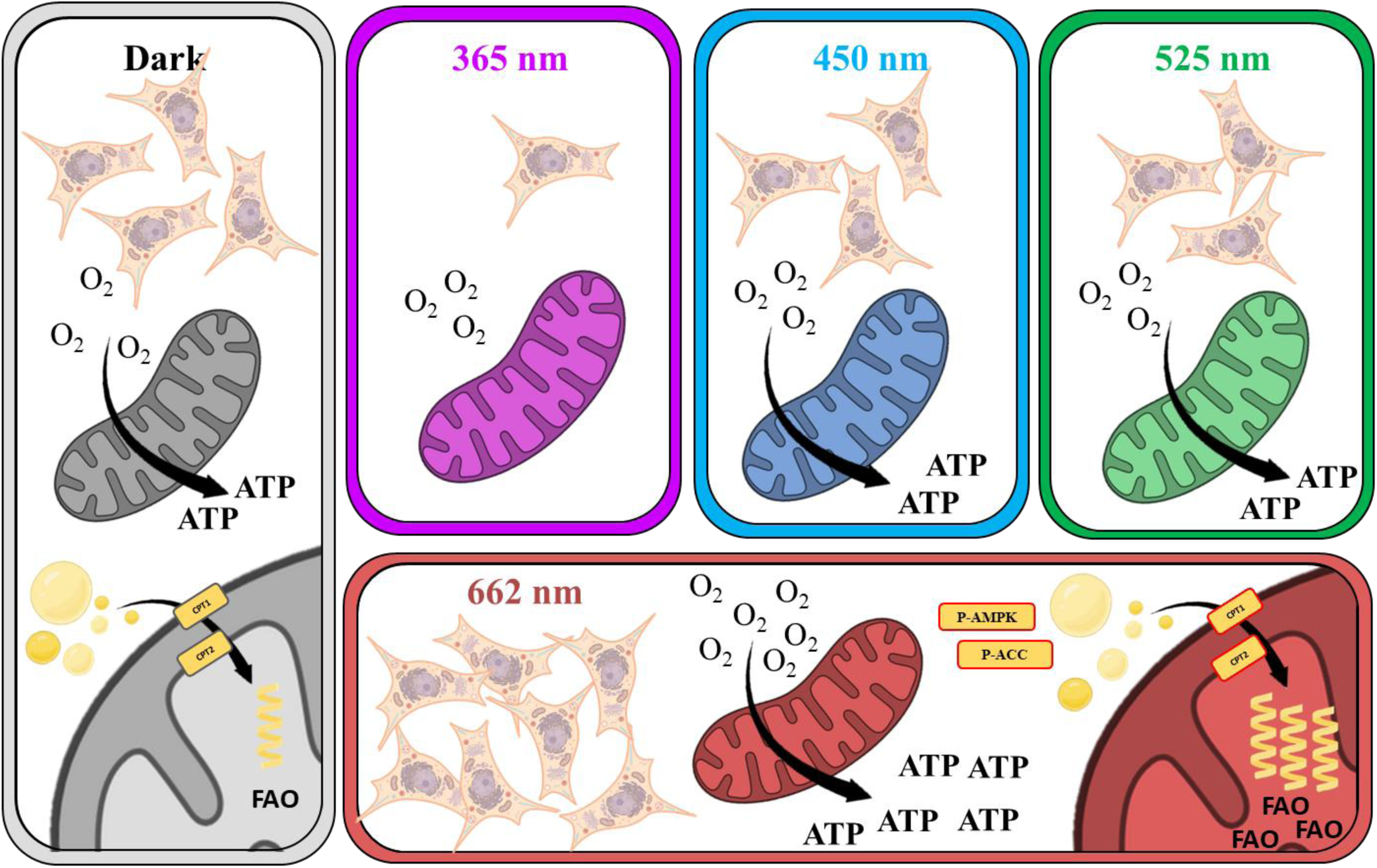
Light at different wavelengths has distinct effects on keratinocyte viability and metabolism. UVA light abrogates metabolic fluxes and cell viability. Blue and green light have no effect on metabolic fluxes, while red light enhanced oxidative phosphorylation by promoting fatty acid oxidation.

## 4. Discussion

Exposure to light is a part of everyday life, and therefore an important aspect to understand scientifically. Historically, light exposure has always been associated with phototoxicity promoted by UVA and UVB light, but more recently visible light ranges such as blue light have also been found to have significant biological effects [15]. In general, most of the biological effects of light described to date are damaging to cells and involve photosensitizing, which generates excited species such as singlet oxygen [5], radicals such as nitric oxide [19] and hydroxyl radicals [20]. This leads to oxidation of important biomolecules including lipids, nucleic acids, and proteins, which can result in unwanted consequences of sunlight exposure such as erythema, photoaging, and cancer.

Given that keratinocytes are the most abundant cell type in the epidermis and that, like all cells, they are rich in endo-photosensitizers, molecules capable of converting light energy into chemical energy by transferring energy or electrons to molecular substrates, we studied here the effects of irradiation with different wavelengths present in sunlight on these cells. We specifically targeted understanding the effects of light on metabolic fluxes, since energy metabolism, and particularly mitochondria, involve high quantities and variety of endo-photosensitizers. Our results confirm that 365 nm UVA exposure at a dose of 36 J/cm^2^ is markedly cytotoxic, leading to both a loss in cell viability (Fig. 1) and abrogated electron transport and oxidative phosphorylation (Fig. 2). Indeed, other groups have found that oxidative imbalance occurs in keratinocytes submitted to UV light [21, 22]. DNA oxidation is one of the targets of photo-oxidation, generating adducts like 8- oxo-7,8-dihydroguanine, which may generate molecular responses such as mutations and, inflammation [23].

Blue light (450 nm) is another wavelength which should be considered when analyzing the cellular effects of sunlight. Indeed, NAD(P)^+^ and NAD(P)H absorb in the blue light range [7] and within membrane compartments can accelerate photosensitized NADH to NAD^+^ conversion [24]. Metabolically, blue light promotes a reduction in maximal respiratory capacity in retina cells [25], as well as increased inflammatory response [26]. Prior studies in retinal cells also find that blue light irradiation at 445 nm correlated with dysregulation of protein levels associated with mitochondrial morphology and dynamics [27]. In epithelial cells, blue light promoted damage to mitochondrial DNA and oxidant production [28]. In keratinocytes, at a dose of 36 J/cm^2^, we find no effect of blue light on viability (Fig. 1) or metabolic fluxes (Fig. 2). This is compatible with prior results from our group showing that blue light affects keratinocyte viability at dose of 60 J/cm^2^ and over [15].

Green light can affect molecules that absorb in this range such as flavins [29, 30]. Indeed, retinal cells have been shown to present decreased viability and electron transport promoted by green light (517 nm, [19, 25]). In adipose cells [31], green light promoted oxidative imbalance related to activation of TRPV1 channels, increasing calcium ion transport. On the other hand, [32] irradiation at 516 nm induces endothelial cell proliferation. Lunova et al., [33] found that irradiation at 502 nm did not promote any effect on inner mitochondrial membrane potentials in hepatic cells, a result that is compatible with our finding that green light did not change basal, ATP-linked, nor maximal mitochondrial electron transport in keratinocytes (Fig. 2), although this wavelength did promote a moderate decrease in cell viability (Fig. 1), through mechanisms that do not involve changes in oxidative phosphorylation.

While damaging effects of light are described in many cells, and expected given spectral properties, we were surprised to see a significant enhancement in basal, ATP-linked and maximal oxygen consumption rates in keratinocytes two hours after being irradiated with red light (660 nm, Fig. 2), an effect that was accompanied by enhanced cell proliferation 48 and 72 hours post-irradiation (Fig. 1). Our results are compatible with those of [34], who found that an epidermal-equivalent cell model and human skin presented increased proliferation after red light treatment, with enhanced mitochondrial oxygen consumption in the cell model. The enhanced oxygen consumption observed occurred in the range of 12-36 J/cm^2^ (Fig. 3), a dosage compatible with expected sunlight exposure [35], and is not related to a thermal effect (Fig. 4). Similarly, mesenchymal stem cells treated with 0.5 a 4 J/cm^2^ red light (630 nm) had significantly increased ATP production and interleukin secretion (IL-6 and IGF-1, [36]). Chu-tan et al., [37] observed an increase of spare respiratory capacity and non-mitochondrial respiration in retinal cells when irradiated with 670 nm red light at different doses. Chu-tan et al., [37] also observed a hormesis effect with low and high doses of light, where higher doses could induce phototoxic effects. This result is compatible with our results in keratinocytes, in which higher doses (150 J/cm^2^) promoted decreased respiration (Fig. 3) and decreased clonogenicity (not shown).

The enhanced oxidative phosphorylation we see in keratinocytes is, therefore, compatible with prior results in the literature. Typically, these effects in prior publications have been associated with photo biomodulation-mediated activation of cytochrome c oxidase in mitochondria directly by red light [38]. However, our results are not compatible with a direct effect of red light on cytochrome c oxidase, for many reasons. First, results are present from two hours (Figs. 2 and 3) to two days (Fig. 5) after irradiation, which would be unexpected for a direct effect of light on enzymatic components. Second, red light increased both basal and maximal oxygen consumption rates (Figs. 2 and 3), while the expected result for cytochrome c oxidase activation in cells with robust reserve respiratory capacity (the difference between basal and maximal oxygen consumption) such as the keratinocytes studied here would be in an increase in maximal oxygen consumption alone. Indeed, the enhanced basal and maximal oxygen consumption rates in the absence of a change in oxygen consumption in the presence of the ATP-synthase inhibitor oligomycin, as observed by us here, suggests an increase in ATP demand and electrons arriving at the electron transport chain from upstream metabolic pathways. These results are in agreement with the observation that cell models lacking cytochrome c oxidase also show the typical photobiomodulation effect of enhancing cell proliferation after red-light irradiation [39].

Consistently, quantification of components of the machinery of oxidative phosphorylation showed no increase that was explanatory of the enhanced oxygen consumption rates observed in cells (Fig. 7). Also, oxygen consumption rates in digitonin-permeabilized cells, where soluble cytosolic components are removed while preserving mitochondrial integrity and function, showed no differences between red light-irradiated and control samples (Fig. 8). Both evidences indicate that red light’s photobiomodulation effects originate from cytosolic metabolic regulation rather than direct modulation of mitochondrial respiratory complexes, challenging the prevailing cytochrome c oxidase absorption paradigm [40]. Using intact cells and monitoring metabolic fluxes dependent on glycolysis (Fig. 9), glutaminase (Fig. 10), and fatty acid oxidation (Fig. 11), we were able to determine that red light irradiation specifically enhanced the latter in keratinocytes. Indeed, etomoxir, an inhibitor of CPT1 and long chain fatty acid entry into mitochondria, specifically decreased respiration in red light-stimulated mitochondria and equalized respiratory rates to those of control cells, confirming that the enhancement is due to fatty acid uptake into mitochondria (Fig. 11), leading to its depletion (Fig. 12). This was correlated with the phosphorylation of AMPK and ACC, regulatory checkpoints for beta oxidation (Fig. 13). While the results involved completely different cell types, our finding that red light induces lipolysis in keratinocytes is compatible with red light treatment (635 nm) effects in rodents, in which it prevented weight gain, hyperlipidemia, and hepatic steatosis promoted by high fat diets [41], our results also correspond with the effect in AMPK phosphorylation in HepG2 cell line observed by Guo et al., 2020 [41].

While mitochondria serve as the central hub of cellular metabolism, their function is also intimately linked to extramitochondrial regulatory networks. Our findings demonstrate that red light photobiomodulation in keratinocytes operates through the activation of AMPK-mediated phosphorylation of ACC to stimulate fatty acid oxidation (FAO), without altering mitochondrial β-oxidation machinery. This reveals a sophisticated light-sensing mechanism wherein: (1) cytoplasmic AMPK serves as the primary photometabolic sensor, (2) ACC inhibition redirects lipid fluxes toward mitochondria, and (3) existing fatty acid oxidation enzymes suffice to enhance respiration. The discovery that red light can trigger lipolysis and elevate basal metabolic rates through this AMPK-ACC-FAO axis provides a mechanistic foundation for understanding phototherapy’s metabolic effects, while opening new questions about how optical signals are transduced into energy homeostasis modulation. These insights position photobiomodulation as a unique tool for probing fundamental connections between light sensing and cellular metabolism.

## 5. Conclusions

Our work shows that keratinocyte viability and metabolism is affected by different wavelengths present in sunlight (see Fig. 13). Intriguingly, while UVA and green light decreased viability, and UVA completely abrogated metabolic fluxes, red light irradiation consistently enhanced oxygen consumption rates in these cells. The effect of red light was dose and cell dependent, long-lasting, and occurred at dosages that are compatible with normal exposure to the sun. Intriguingly, the effects of red light in keratinocytes were not related to direct changes in mitochondrial metabolism, but instead to a specific enhancement of fatty oxidation in irradiated cells via phosphorylation of AMPK and ACC proteins. This new and interesting effect of light may explain many of its effects in animal models, and expands the possibilities of use of red light in physiological cell modulations and therapeutics.

## Abbreviations

2DG: 2-deoxy-glucose;
III-UQCRC: Q cytochrome C oxidoreductase;
I- NDUFBS: NADH dehydrogenase;
IV-COX II: cytochrome C oxidase;
ADP: Adenosine Diphosphate;
ACC: Acetyl-CoA carboxylase;
AMP: Adenosine Monophosphate;
AMPK: AMP-activating protein kinase;
ATP: Adenosine Triphosphate;
CCCP: Carbonyl cyanide m-chlorophenyl hydrazone;
CPT1: Carnitine palmitoyl-transferase I;
CPT2: Carnitine palmitoyl-transferase II;
CV: crystal violet;
CVS: Cristal Violet Staining;
DMSO: dimethyl sulfoxide;
ECAR: Extra-Cellular Acidification Rates;
ETFA: Electron-transfer- flavoprotein;
FAO: Fatty acid oxidation;
FBS: fetal bovine serum;
FFA: Free Fatty Acids;
MTT: 3-(4,5-dimethylthiazol-2-yl)-2,5-diphenyltetrazolium bromide;
NEFA: Non-Esterified Free Fatty Acids;
NR: Neutral Red;
NRS: neutral red staining;
OCR: Oxygen Consumption Rates;
P-ACC: Acetyl-CoA carboxylase phosphorylated;
P-AMPK: AMP-activating protein kinase phosphorylated;
PBS: phosphate buffer saline;
PDH: Pyruvate dehydrogenase;
R/AA: rotenone and antimycin;
RCR: Respiratory control ratios;
SDHB: Succinate dehydrogenase;
V-ATP5A: ATP Synthase;
UVB: Ultraviolet B;
UVA: Ultraviolet A.

## Author contributions

All authors participated in experimental design, data analysis, reading and approval of the final manuscript. MAHL and CCCS conducted experiments.

## Supporting information

supl mat

## Acknowledgments

This work was supported mainly by the Fundação de Amparo à Pesquisa do Estado de São Paulo (FAPESP) Supported by FAPESP grant numbers 20/06970-5 and 2021/08521- 6, Conselho Nacional de Pesquisa e Desenvolvimento (CNPq), Coordenação de Aperfeiçoamento de Pessoal de Nível Superior (CAPES) line 01 and the Centro de Pesquisa, Inovação e Difusão de Processos Redox em Biomedicina - CEPID Redoxoma, grant 2013/07937-8. MAHL is supported by a FAPESP fellowship. We thank Sirley Mendes de Oliveira, Helena C. Junqueira, and Julian D. Serna for the excellent technical assistance and data discussion.

