## Supplementary material for "Mitochondrial Fatty Acid Oxidation is Stimulated by Red Light Irradiation": supl mat

**Figure Legends**

**Figure S1. Spectral distributions of LEDs used for irradiation treatments:** UVA (365 nm), blue light (450 nm), green light (517 nm), and red light (660 nm). Irradiance was measured as described in the methods section.

**Figure S2. Mitochondrial respiration is differentially modulated by different light wavelengths.** Cell oxygen consumption rates (OCR) and Extracellular Acidification Rates (ECAR) were quantified after 2 hours irradiation at 36 $J/cm^2$ with UVA (365 nm), blue light (450 nm), green light (517 nm), or red light (660 nm). OCRs/ECARs were measured as described in Methods under basal conditions, followed by injection of oligomycin (oligo, 1 μM), CCCP (1 μM), and antimycin A plus rotenone (AA/Rot, 1 μM each). Results are means ± SD of three independent experiments; ns = > 0.1. One-way ANOVA followed by Dunnett.

**Figure S3. Moderate doses of red light (660 nm) increase mitochondrial respiration (continuation).** OCRs and ECAR were quantified under similar conditions to Fig. 2 after 2 hours irradiation at 6, 12, 36 and 150 $J/cm^2$ with red light. Results are expressed as mean ± SD of three independent experiments; ns = > 0.1, One-way ANOVA followed by Dunnett.

**Figure S4. No changes in proteins related to fatty acid oxidation upon treated with Red light.** Western blot analysis of protein levels was conducted in cells 2 hours post exposure to red light at different doses (0, 12, 36 $J/cm^2$). Extracts were collected and protein quantified by BCA. Carnitine palmitoyl transferases (CPT1/CPT2) (A, B), Electron-transfer-flavoprotein (ETFA) (C) and Pyruvate dehydrogenase (PDH) (D) where quantified. Results are expressed as means ± SD of four independent experiments; ns = > 0.1, One-way ANOVA followed by Dunnett.

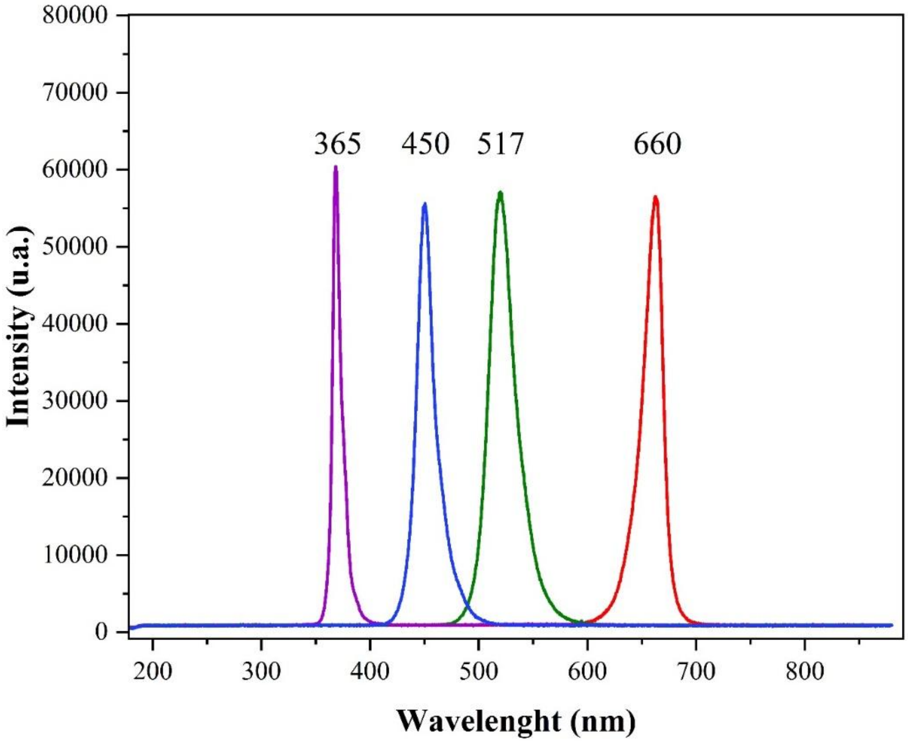


Herrera et al., Supplementary Figure S1

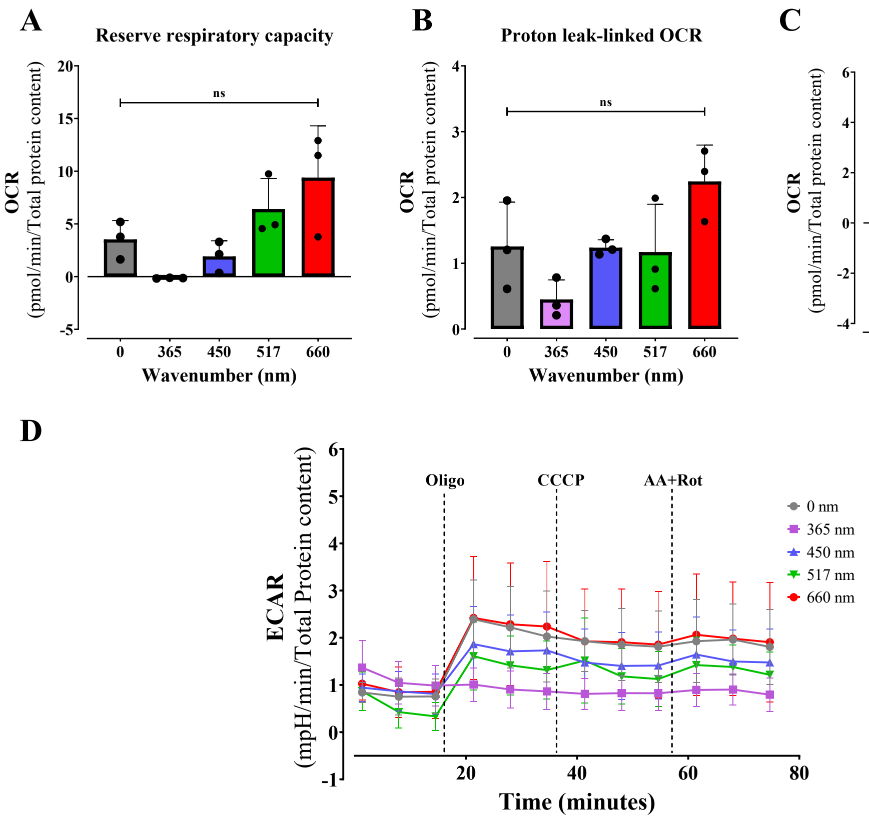


Herrera et al., Supplementary Figure S2

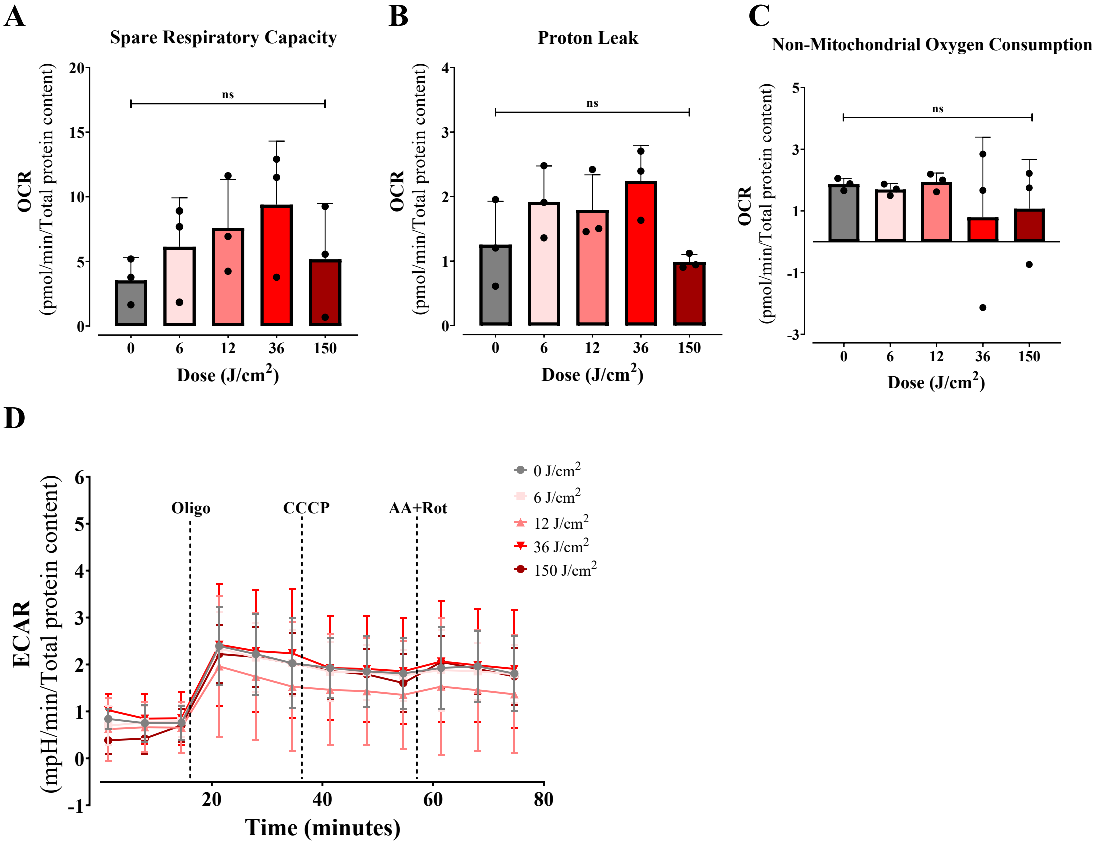


Herrera et al., Supplementary Figure S3

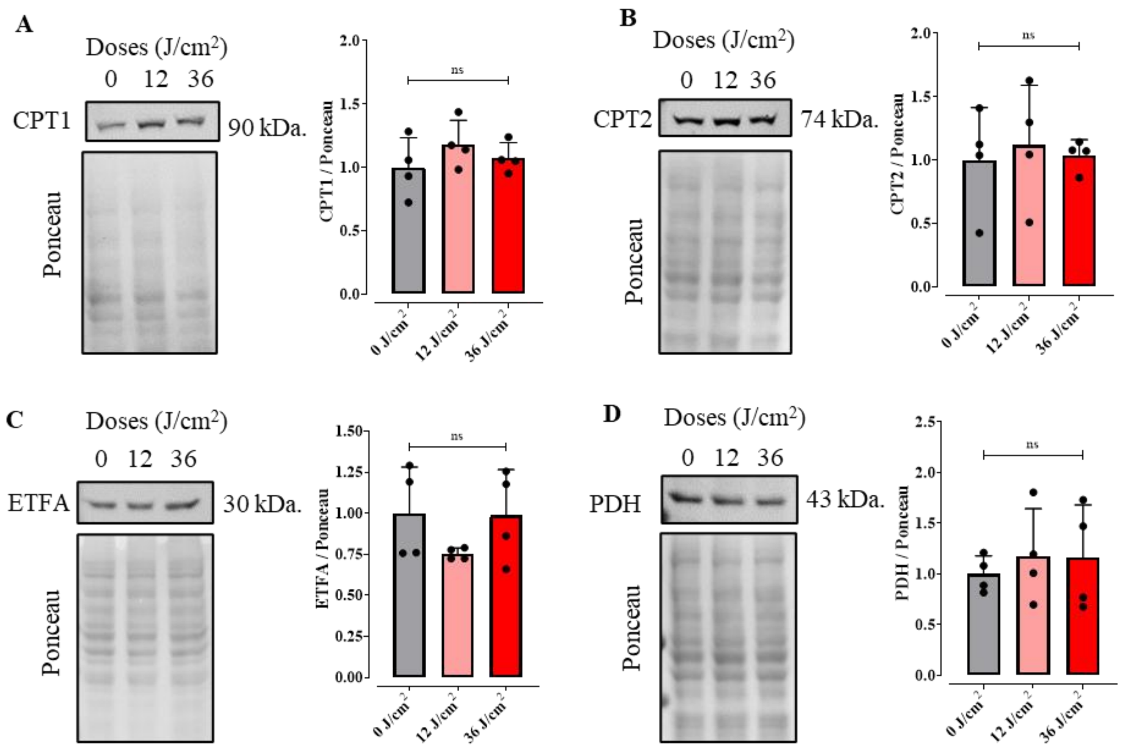


Herrera et al., Supplementary Figure S4

**Supplementary tables**

**Table S1**. LED specifications and photons from each irradiator used for UVA (365 nm), Blue light (450 nm), Green light (517 nm) and Red light (660 nm) with a dose of 36 J/cm$^2$. Energy was calculated using the formula E= hc/λ, where "h" is the Planck's constant, "c" is the speed of light in vacuum and λ is the wavelength.

| Wavelength (m) | Intensity (mW) | Energy (J) | Number of photons / s | Dose (J/cm$^2$) | Area (cm$^2$) | Time (s) | Total number of photons |
|---|---|---|---|---|---|---|---|
| 3,65 x $10^{-7}$ | 2,16 | 5,45 x $10^{-19}$ | 3,96 x $10^{-18}$ | 36 | 0,283 | 4716,67 | 1,87 x $10^{-22}$ |
| 4,50 x $10^{-7}$ | 16,70 | 4,42 x $10^{-19}$ | 3,78 x $10^{-19}$ | 36 | 0,49 | 1056,29 | 3,99 x $10^{-22}$ |
| 5,17 x $10^{-7}$ | 13,00 | 3,85 x $10^{-19}$ | 3,38 x $10^{-19}$ | 36 | 0,49 | 1356,92 | 4,59 x $10^{-22}$ |
| 6,60 x $10^{-7}$ | 12,20 | 3,01 x $10^{-19}$ | 4,05 x $10^{-19}$ | 36 | 0,49 | 1445,90 | 5,85 x $10^{-22}$ |

**Table S2.** LED specifications and photons of the Ethik irradiator used for red light treatments (660 nm) with the different doses (6, 12, 36 and 150 J/cm$^2$). Energy was calculated using the formula of E= hc/λ, where "h" is the Planck's constant, "c" is the speed of light in vacuum and λ is the wavelength.

| Wavelength (m) | Intensity (mW) | Energy (J) | Number of photons / s | Dose (J/cm$^2$) | Area (cm$^2$) | Time (s) | Total number of photons |
|---|---|---|---|---|---|---|---|
| 6,60 x $10^{-7}$ | 12,20 | 3,01 x $10^{-19}$ | 4,05 x $10^{-19}$ | 6 | 0, 49 | 240,98 | 9,76 x $10^{-21}$ |
| 6,60 x $10^{-7}$ | 12,20 | 3,01 x $10^{-19}$ | 4,05 x $10^{-19}$ | 12 | 0,49 | 481,96 | 1,95 x $10^{-22}$ |
| 6,60 x $10^{-7}$ | 12,20 | 3,01 x $10^{-19}$ | 4,05 x $10^{-19}$ | 36 | 0,49 | 1445,90 | 5,85 x $10^{-22}$ |
| 6,60 x $10^{-7}$ | 12,20 | 3,01 x $10^{-19}$ | 4,05 x $10^{-19}$ | 150 | 0,49 | 6024,59 | 2,44 x $10^{-23}$ |
